# Quantifying dynamic pro-inflammatory gene expression and heterogeneity in single macrophage cells

**DOI:** 10.1101/2023.05.19.541501

**Authors:** Beverly Naigles, Avaneesh V Narla, Jan Soroczynski, Lev S Tsimring, Nan Hao

## Abstract

Macrophages must respond appropriately to pathogens and other pro-inflammatory stimuli in order to perform their roles in fighting infection. One way in which inflammatory stimuli can vary is in their dynamics – that is, the amplitude and duration of stimulus experienced by the cell. In this study, we performed long-term live cell imaging in a microfluidic device to investigate how the pro-inflammatory genes IRF1, CXCL10, and CXCL9 respond to dynamic interferon-gamma (IFNγ) stimulation. We found that IRF1 responds to low concentration or short duration IFNγ stimulation, whereas CXCL10 and CXCL9 require longer or higher-concentration stimulation to be expressed. We also investigated the heterogeneity in the expression of each gene and found that CXCL10 and CXCL9 have substantial cell-to-cell variability. In particular, the expression of CXCL10 appears to be largely stochastic with a subpopulation of non-responding cells across all the stimulation conditions tested. We developed both deterministic and stochastic models for the expression of each gene. Our modeling analysis revealed that the heterogeneity in CXCL10 can be attributed to a slow chromatin-opening step that is on a similar timescale to that of adaptation of the upstream signal. In this way, CXCL10 expression in individual cells can remain stochastic in response to each pulse of repeated simulations, as validated by experiments. Together, we conclude that pro-inflammatory genes in the same signaling pathway can respond to dynamic IFNγ stimulus with very different response features and that upstream signal adaptation can contribute to shaping the features of heterogeneous gene expression.

## Introduction

Cells respond to external signals by initiating gene expression programs to elicit appropriate physiological responses. However, even within a clonal population, there can be significant variability in gene expression among individual cells (1–4). Studies have shown that several mechanisms, including environmental fluctuations, epigenetic regulation, and the inherent stochasticity of biochemical reactions can potentially contribute to this heterogeneity (5–9). Importantly, cell-to-cell variability in gene expression can lead to functional consequences for development, disease progression, and response to therapy (10, 11). For example, some cancer cells in a tumor can be more resistant to chemotherapy than others due to differences in gene expression involved in drug metabolism or DNA repair (12–14). Similarly, during hematopoiesis, the high expression heterogeneity of the stem cell marker Sca-1 leads to different fate decisions toward erythroid or myeloid lineages (15). In this study, we focus on heterogeneity in the immune response and investigate the dynamics of gene expression in single macrophage cells.

Macrophages are innate immune cells that perform a diverse range of functions in the body and adapt their functional response to their local signaling environment. Macrophage phenotypes exist along a continuous spectrum and single cells dynamically shift between states (16), but here we focus on the pro-inflammatory M1 phenotype. This phenotype, wherein macrophages play anti-microbial roles and promote inflammatory immune responses, occurs *in vivo* in response to bacterial or viral infections and is modeled *in vitro* by exposure to lipopolysaccharide (LPS) or interferon-gamma (IFNγ) (17).

Macrophages are highly heterogeneous cells and acrophage heterogeneity has been studied *in vitro* both before and after infection (18–20). In mouse bone marrow-derived macrophages infected with *Salmonella enterica*, infected macrophages adopt different gene expression states with varying levels of pro-inflammatory gene expression (21). In diseases such as tuberculosis, variability in how macrophages respond to infection leads to dramatic differences in clinical outcome (22, 23). This macrophage heterogeneity is also seen *in vivo*, where macrophages have diverse functions and gene expression patterns in many organs, including the lung (24), brain (25), and peritoneum (26), as well as in response to inflammation (27).

Single-cell RNA sequencing (scRNAseq) has uncovered gene expression heterogeneity among individual macrophages and other myeloid cells exposed to pro-inflammatory signals such as LPS and IFNγ *in vitro*. Such experiments eliminate the element of bacterial heterogeneity and have shown that heterogeneity is present in aspects of immune responses other than direct interaction with a pathogen (28–31). However, these scRNAseq studies cannot reveal how these single-cell transcriptional patterns vary in time in response to pro-inflammatory stimuli. Bulk studies show that pro-inflammatory gene expression is highly temporally regulated, with sets of earlier primary response genes and later secondary response genes that each share certain elements of their epigenetic and transcriptional regulation (32, 33). However, the intersection of heterogeneity and gene expression timing remains poorly understood.

The main system in which pro-inflammatory gene expression in macrophages has been studied at the single-cell level over time is in genes induced by the transcription factor NFkB, which translocates to the nucleus upon stimulation with pathogens or pathogen analogs. NFkB activates different gene expression programs based on its residence time dynamics in the nucleus, and the residence time dynamics depend on the identity of the upstream signal and can vary between clonal cells (34–37). Varied transcription factor dynamics leading to differential gene expression and cellular outcome have also been seen in other systems such as the yeast Msn2 system and p53 expression in the MCF7 breast cancer cell line (38–40).

These studies used quantitative analysis of single-cell time traces coupled with mathematical modeling to uncover the gene expression networks and mechanisms that encode and decode complex environmental signals, since bulk measurements of dynamical behavior can distort individual patterns due to averaging over different single cells (39). Observing gene expression output in response to dynamic upstream signals can reveal elements of network structure, whether the dynamic upstream signal is natural (e.g. ERK signaling in response to EGF vs NGF (41)) and/or is exogenously applied (38, 42, 43). Mathematical modeling can rule in or out possible underlying network motifs and mechanisms of epigenetic regulation of gene expression, as well as suggesting perturbations that will change the dynamical behavior of the system, which can then be tested (42, 44).

Macrophage signal processing and heterogeneity is essential for properly modulating immune responses both at initial pathogen recognition and within the later stages of the innate and adaptive immune response; however, signal processing in the later stages of the innate and adaptive immune response is poorly studied. To this end, we focused on the cytokine interferon gamma (IFNγ), which polarizes macrophages to an M1 phenotype but is secreted by other immune cells rather than directly associated with a pathogen. Macrophages encounter IFNγ in various temporal patterns during the immune response since it is secreted by natural killer cells during the innate immune response and CD4 Th1 cells during the adaptive immune response (45, 46). IFNγ binds to the interferon-gamma receptor and signals through JAK-STAT signaling, leading to STAT1 homodimerization, phosphorylation, and translocation into the nucleus where it acts as a transcription factor for the downstream gene expression program (45). Mis-regulation of IFNγ signaling is pervasive in disease, from too little IFNγ leading to poor infection response, to too much IFNγ leading to excessive inflammation and autoimmune diseases, to dysregulation of IFNγ being important in tumor immunology (47–49).

In this study, we investigate the single-cell expression dynamics of the genes IRF1, CXCL10, and CXCL9, which are all induced by STAT1 in response to IFNγ. IRF1 and CXCL10 are representative primary response genes with different kinetics (early for IRF1, later for CXCL10), and CXCL9 is a secondary response gene (50). IRF1 encodes a transcription factor important for expression of many other important pro-inflammatory genes (50–53). CXCL10 and CXCL9 encode chemokines that recruit T cells and other cells expressing the CXC chemokine receptor 3 (CXCR3) to the site of inflammation and have been implicated in a number of disease conditions (54–56). As an example, CXCL10 expression is important early in SARS-CoV-2 infection to create an anti-viral environment, but later in infection high CXCL10 and CXCL9 contribute to the cytokine storm that leads to severe disease (57, 58).

## Results

### Quadruple-reporter macrophage cell lines to study IFNγ-induced gene expression

We used CRISPR/Cas9 genome editing to create a RAW264.7 mouse macrophage-like cell line that expressed endogenous fluorescent reporters for each of IRF1, CXCL10, and CXCL9 in the same single cells, as well as a nuclear marker for use in image analysis (Fig. 1A and Methods). RAW264.7 cells have been used as a model for much of the work studying NFkB signaling dynamics in macrophages (19, 34, 59–62), gene regulation in pro-inflammatory macrophages (63–65), and *in vitro* macrophage models for mycobacterial infection (66, 67). The cell line created in our study is heterozygous for the SYFP2 tag at the IRF1 locus, and homozygous for both the mCerulean knock-in at the CXCL10 locus and the mCherry knock-in at the CXCL9 locus. For CXCL10 and CXCL9 the fluorescent protein DNA sequence is connected to the endogenous chemokine DNA sequence by a T2A translational skip site, ensuring that the endogenous chemokine can be secreted normally and the NLS-fluorescent reporter protein is retained in the nucleus where it can be measured (Fig. 1A). A list of all cell lines created for this work can be found in Supplementary Table 4.

**Figure 1.**
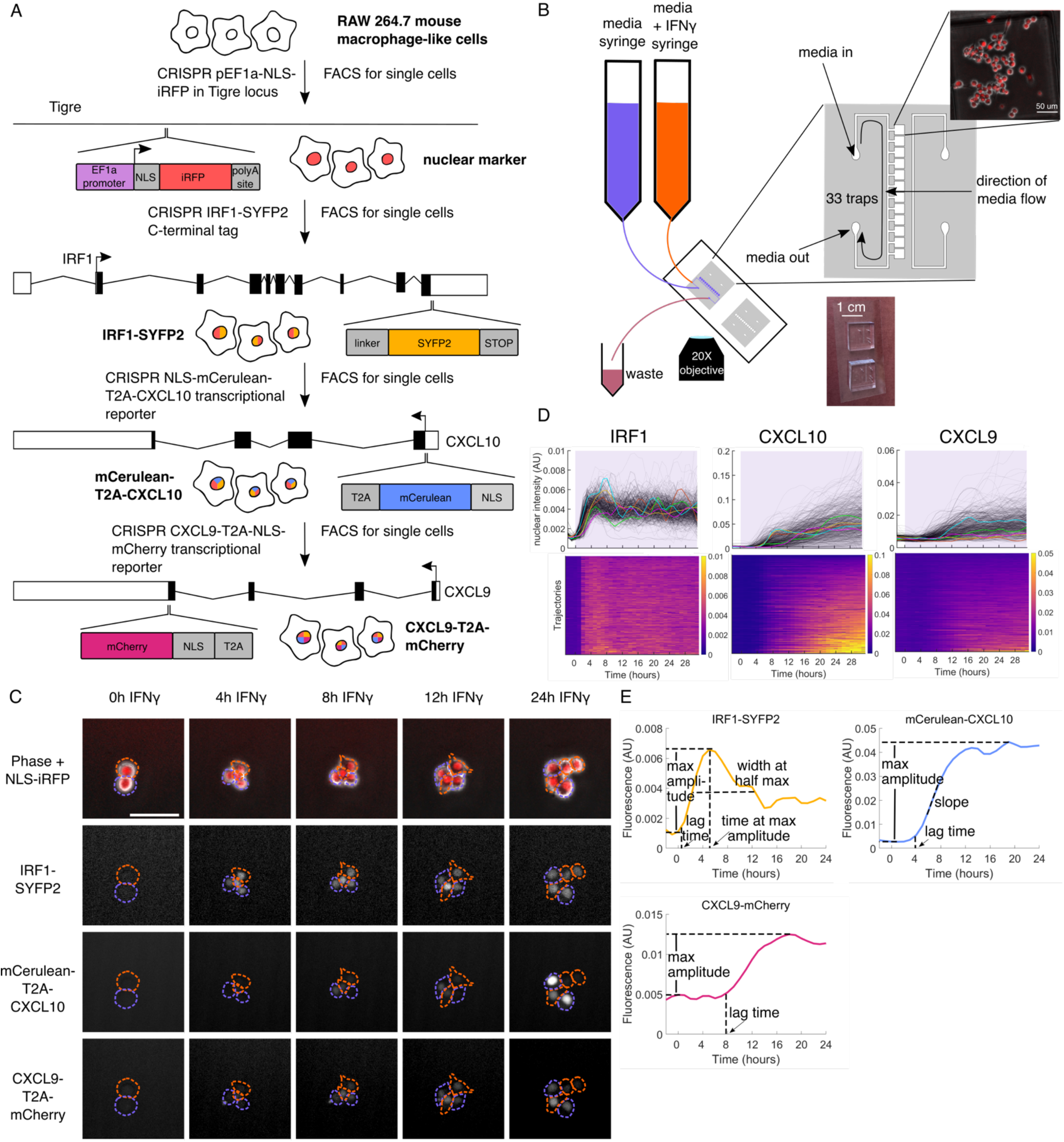
Endogenous CRISPR knock-in cells for measuring IFNγ-induced gene expression in single macrophage cells over time. A. Schematic of cell line construction. B. Diagram of microfluidic chip and setup used in these experiments. C. Sample images of cells experiencing IFNγ stimulus. Single cells are outlined over time, with the same color outline showing the same cell and its children. Scale bar on top left image represents 50 μm. D. IRF1, CXCL10, and CXCL9 gene expression responses to 10 ng/mL IFNγ in a 24-well plate. In the top row, each grey line is a cell, with five traces colored as examples. The bottom row shows the same data as heatmaps with each row representing one cell. The heatmaps are sorted by the maximum fluorescence value for each cell for each gene, resulting in the sort order being different for each of the three genes. IFNγ is added at time 0. Color in the heatmaps represents gene expression fluorescence. E. Schematic of sample traces for IRF1, CXCL10, and CXCL9 showing the features extracted from each trace.

We then exposed these cells to IFNγ in either a 24-well plate or a microfluidic device for thirty-one hours taking fluorescence images every hour. We used an image-processing pipeline to extract fluorescence measurements for each gene over time in single cells (Methods). The microfluidic device allows us to expose the cells to time-variant stimulus patterns and have media turnover in less than 15 minutes (Fig. 1B) (68). Sample single-cell traces are shown in Fig. 1D, and a sample field of view in Fig. 1E, where we can see substantial heterogeneity in CXCL10 and CXCL9 expression between cells.

Looking at these traces, we can extract several features that together describe the response for each gene (Fig. 1E). We are also interested in how these features correlate with each other and change in response to dynamic stimulus, which we will describe below. For each gene we measure the lag time before protein fluorescence can be measured, as well as the maximum expression amplitude (Fig. S1 J, K, O, P, T, U). Additionally, for CXCL10 we measure the slope of the response, and for IRF1 the width of the peak and the time at the maximum value (Fig. S1 L, M, Q).

To confirm that IFNγ does not induce global changes in gene expression, we measured nuclear marker expression upon IFNγ stimulus and saw that its levels remain constant (Fig. S1B). We also confirmed that the pre-stimulus expression levels of IRF1, CXCL10, and CXCL9 do not correlate with their eventual maximum fluorescence levels (Fig. S1 C-F). To determine if there is correlation between these genes on a single-cell level, we correlated maximal expression levels of IRF1, CXCL10, and CXCL9 and see no correlation between IRF1 and either CXCL10 or CXCL9, and a weak positive correlation between CXCL10 and CXCL9 (Fig. S1A, G-I).

When we cross-correlate features other than amplitude of different genes in the same cells we see no strong correlations (Fig. S1 N, R, S, V, W). Based on these data, we will treat each gene independently in our analysis.

### IRF1, CXCL10, and CXCL9 are expressed differently in response to stimuli of varying concentrations or durations

To investigate how these gene expression features change in response to dynamic stimuli, we first exposed the cells to constant IFNγ stimulation of different concentrations. IRF1 shows fast and uniform expression in all cells, with the cells responding even to 0.1 ng/mL IFNγ and saturating at 3 ng/mL IFNγ. (Fig. 2A and D). When tissue IFNγ levels have been measured after infection, they have ranged from 0.1 ng/mL to 10 ng/mL, showing that this is a physiologically relevant range(69, 70). The width and timing of the IRF1 peak do not vary with the concentration of IFNγ and at all doses we see that the majority of cells reach their maximal expression around four to six hours after onset of stimulation and have a peak width around nine hours, with a minority of cells peaking later (Fig. S2C and D). At a single-cell level, neither peak time nor width correlates with expression amplitude. However, we note that peak width correlates positively with peak time. This is explained by the fact that later peaks are due to IRF1 expression staying high after the initial rise, resulting in a wider peak (Fig. S2 H-J).

**Figure 2.**
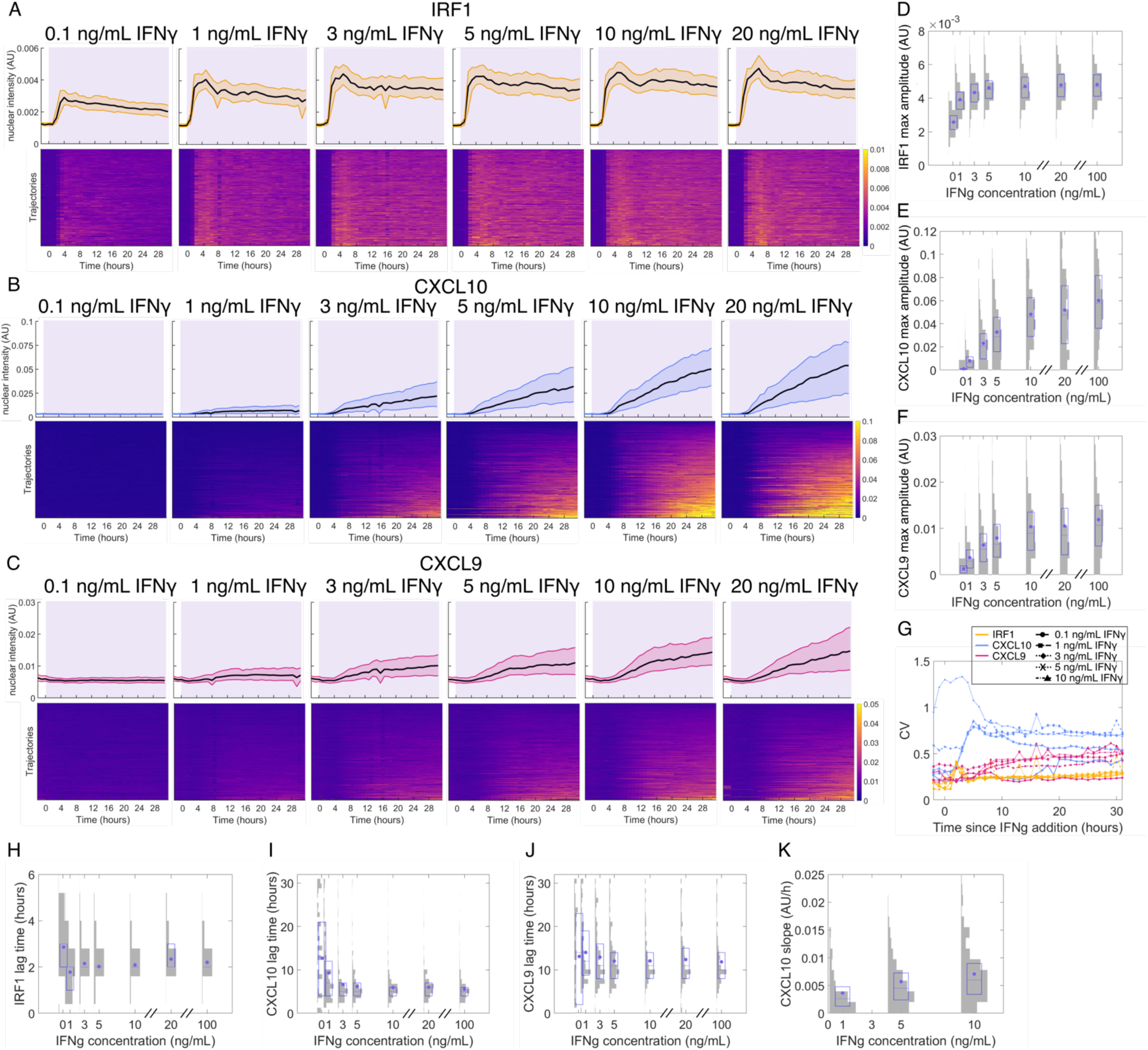
IRF1, CXCL10, and CXCL9 response to varying concentrations of IFNγ stimulation. A-C. Gene expression responses for 0.1-20 ng/mL IFNγ in a 24-well plate with IFNγ added at 0 h. The top row of plots for each gene has a black line at the median and shading to the 25^th^ and 75^th^ percentiles, and each row in the heatmaps in the bottom row shows data for each individual cell. Each heatmap is sorted vertically by single-cell maximum value for that specific gene. Purple shading indicates when the cells are exposed to IFNγ. Color in the heatmaps represents gene expression amplitude. A is IRF1, B is CXCL10, C is CXCL9. D-F. Histograms overlaid with box plots showing maximum IRF1 (D), CXCL10 (E) or CXCL9 (F) expression in single cells for each IFNγ concentration. G. Coefficient of variation of expression at each timepoint (see methods) for all genes across IFNγ concentrations. H-J. Histograms overlaid with box plots showing the lag time for IRF1 (H), CXCL10 (I), and CXCL9 (J) in single cells for each IFNγ concentration. K. Histograms overlaid with box plots showing the CXCL10 slope in single cells for each IFNγ dose. For all box plots, the purple dot is the mean, the purple center line is the median, and the purple box is the 25^th^-75^th^ percentile. The grey shading shows the histogram distribution among single cells for each condition.

In contrast, CXCL10 shows slower and more heterogeneous expression, which saturates at 10 ng/mL IFNγ and has a sharp increase in amplitude from 0.1 ng/mL to 10 ng/mL (Fig. 2B and E). CXCL10 lag time does not vary with stimulus concentration and has a narrow and symmetric distribution (Fig. 2I) indicating that cells begin to express CXCL10 at a defined time regardless of stimulus concentration. CXCL10 lag time also does not correlate with gene expression amplitude on a single cell level (Fig. S2K). In contrast, CXCL10 slope increases with stimulus dose up to 10 ng/mL on a population level, and in single cells the slope correlates positively with CXCL10 expression amplitude (Figs. 2K and S2L). Intriguingly, we observe that the proportion of cells that express high levels of CXCL10 increases with increasing IFNγ concentration, but at all concentrations there is always a fraction of cells with very low or no CXCL10 expression (Fig. S2N). The proportion of these non-responding cells remains unchanged even when IFNγ concentration is increased beyond saturation (10-100 ng/mL) (Figs. 2B and S2B and N). Similarly to CXCL10, CXCL9 expression increases nonlinearly in response to IFNγ doses from 0.1 ng/mL up to 10 ng/mL (Fig. 2F), shows substantial heterogeneity (Fig. 2C), and has a lag time that remains unchanged across IFNγ doses and is uncorrelated with expression amplitude in single cells (Figs. 2J and S2M).

We next exposed the cells to 1-hour, 4-hour, 8-hour, or constant durations of 10 ng/mL IFNγ in a microfluidic device. We chose 10 ng/mL because it is the saturating concentration in our system. Cells show a homogeneous IRF1 peak with a similar amplitude and lag time across all input durations (Fig. 3A, D, and H), while the width of the peak increases with input duration until it saturates between 8 and 24 hours (Fig. 3A and K). For CXCL10, increasing IFNγ input duration leads nonlinearly to increased gene expression, with 1 hour of IFNγ inducing very low expression and expression amplitude saturating between 4 hours and 8 hours of IFNγ treatment (Fig. 3B and E). The CXCL10 slope is also lower with shorter IFNγ duration but saturates at around 4 hours of stimulation (Fig. S3B). The CXCL10 lag time is similar for all IFNγ durations of four hours or longer, while for one hour of stimulation the cells that do express CXCL10 have a very short lag time, showing that this 1-hour stimulation only induces the earliest cells (Fig. 3I). Increasing IFNγ duration also increases the proportion of cells that strongly express CXCL10, but there remains a fraction of cells with very low CXCL10 expression even in response to constant stimulation (Figs. 3B and S3G). CXCL9 expression also increases with input duration but does not saturate by 24h; the cells continue to produce CXCL9 for as long as stimulus is provided (Fig. 3C, F).

**Figure 3.**
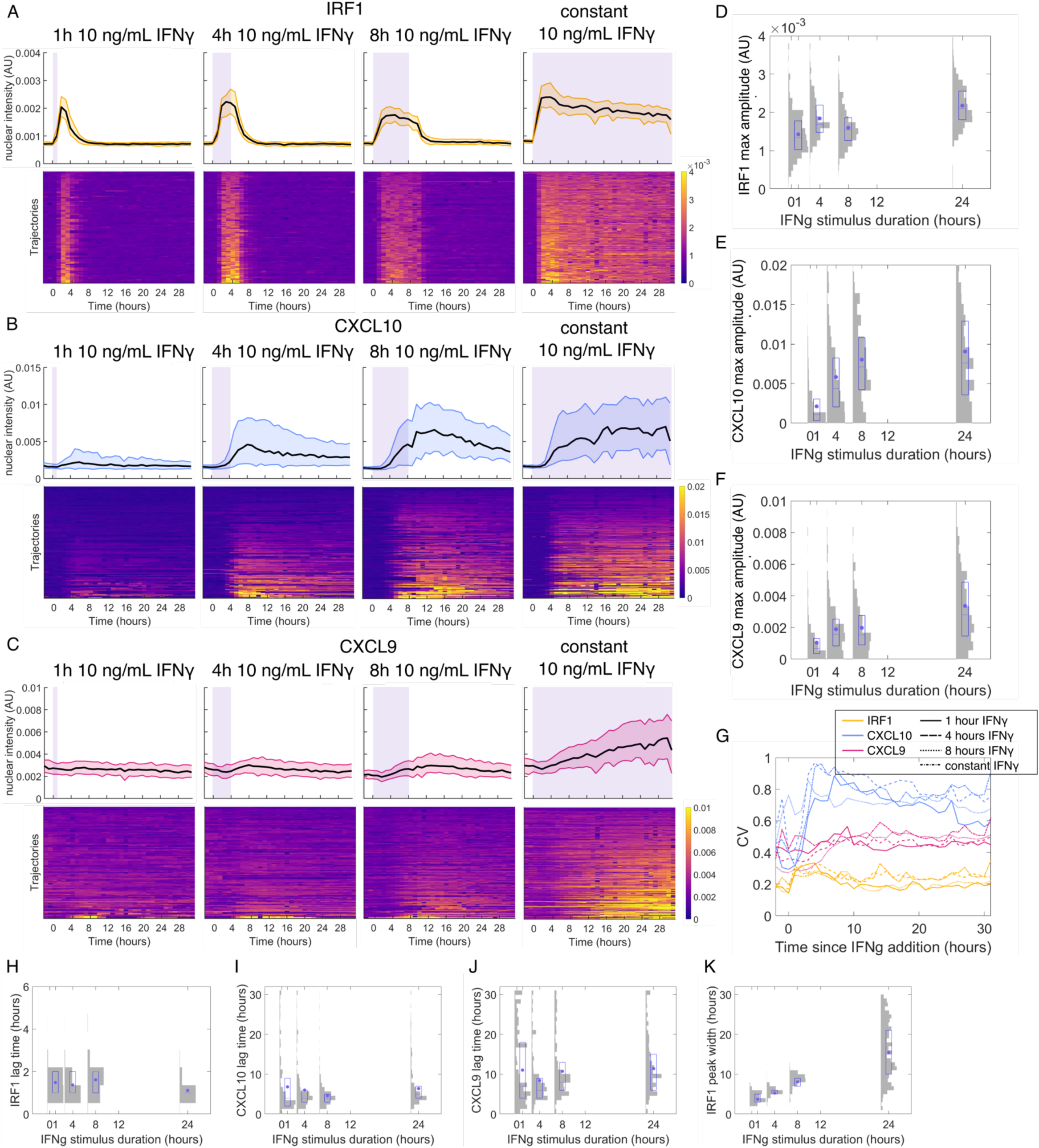
IRF1, CXCL10, and CXCL9 response to varying durations of IFNγ stimulation. A-C. Gene expression responses for 1 h, 4 h, 8 h, and constant 10 ng/mL IFNγ stimulation in a microfluidic device with IFNγ added at time 0. The top row of plots for each gene has a black line at the median and shading to the 25^th^ and 75^th^ percentiles, and each row in the heatmap represents one cell. Each heatmap is sorted by single-cell maximum value for that specific gene. Purple shading indicates the time during which the cells are exposed to IFNγ. Color in the heatmap corresponds to gene expression amplitude. A is IRF1, B is CXCL10, C is CXCL9. D-F. Histograms overlaid with box plots showing maximum IRF1 (D), CXCL10 (E) or CXCL9 (F) expression in single cells for each IFNγ duration. G. Coefficient of variation of expression at each timepoint (see methods) for all genes across IFNγ durations. H-J. Histograms overlaid with box plots showing the lag time for IRF1 (H), CXCL10 (I), and CXCL9 (J) in single cells for each IFNγ duration. K. Histograms overlaid with box plots showing the IRF1 peak width in single cells for each IFNγ dose. For all histograms overlaid with box plots, the purple dot is the mean, the center purple line is median, and the purple box is the 25^th^-75^th^ percentile. Grey shading shows the histogram distribution among single cells for each condition.

Taken together, these results demonstrate that the IRF1expression is fast and uniform among cells, whereas CXCL10 expression is slower and highly heterogeneous, and CXCL9 expression is even slower than CXCL10 and also heterogeneous. IRF1 responds strongly to low-concentration or short-duration IFNγ stimulation, while CXCL10 and CXCL9 act as high-pass filters to filter out these lower stimuli and respond strongly only to higher-concentration or longer stimulation. We see that lag time is invariant to dynamic stimulation (as would be expected as cells have no knowledge of future events) and that the lag time distributions for IRF1 and CXCL10 are narrow while the distributions for CXCL9 are wider (Fig. 2 H-J). This shows that heterogeneity in CXCL10 expression comes from expression amplitude rather than timing, but that CXCL9 expression also varies in timing. Other than amplitude, the features that vary between cells or with dynamic input are CXCL10 slope, which increases with expression amplitude, and IRF1 peak width, which increases with IFNγ duration. We see no identifiable spatial pattern in the CXCL10 and CXCL9 heterogeneity.

In addition, for CXCL10, increasing either input amplitude or duration can increase the proportion of cells with high expression levels of the gene; however, even upon constant stimulation with saturating concentrations of IFNγ, there remains a substantial fraction of cells with low-to-no expression. To quantify cell-to-cell variation in gene expression, we calculated the coefficient of variation (CV) for each gene (see Methods) and observed that CXCL10 has the highest CVs and IRF1 has the lowest under all the stimulation conditions tested (Figs. 2G and 3G). The CV for all genes varies minimally across input concentrations and durations, despite the fact that the mean expression level of CXCL10 is altered by up to 6-fold (Figs. 2G and 3G).

### Computational modeling of gene expression responses to dynamic stimuli

Two features of our dynamic stimulation data were particularly striking to us: the different ways in which each gene either filtered out or responded to IFNγ stimulus of low amplitude or short duration, and the persistence of cells with low-to-no CXCL10 expression in all conditions. To investigate the possible mechanisms of these phenomena, we constructed deterministic and stochastic models of this signaling and gene expression system. We began with an ordinary differential equation (ODE) model that describes chromatin opening, transcription initiation, mRNA synthesis, mRNA translation, and fluorescent protein maturation, as well as degradation of the mRNA and protein (Fig. 4A). We fit this model to the mean protein time courses in response to IFNγ stimulations of all amplitudes and durations for each of IRF1, CXCL10, and CXCL9 (Fig. 4C, Supp Table 1 for best fit parameters, Methods for detailed description of modeling). For CXCL10 we fit only to the first twelve hours of the plate (concentration) experiments because we see that after twelve hours the CXCL10 response in a microfluidic device stops increasing, but in the plate it continues to increase. This is likely because the cells are secreting a paracrine effector that leads to increased CXCL10 expression over time in the plate experiments but is washed out in the microfluidic experiments. As we are not trying to model paracrine effectors, we chose to ignore this region in the plate experiments. When we compare the parameters and look at the intermediate model states, we see that in our model chromatin opening and transcription initiation are much slower for CXCL10 than for IRF1, and slower for CXCL9 than CXCL10 (Supp. Table 1 and Fig. S4A).

**Figure 4.**
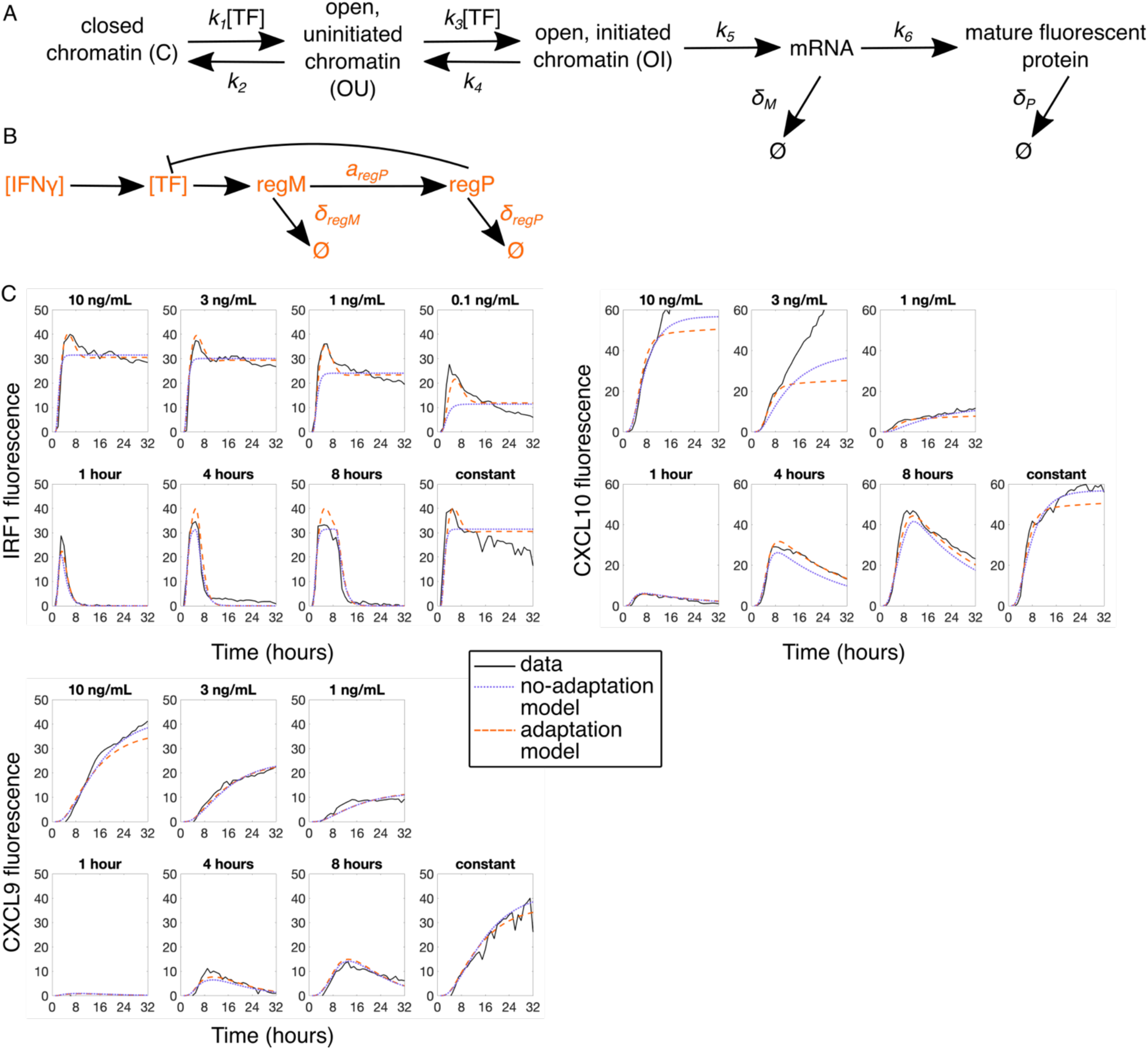
Deterministic modeling of IFNγ-induced gene expression. A. Schematic diagram of the gene expression model architecture with 3-state chromatin dynamics and gene transcription and translation for all three genes of interest (IRF1, CXCL10, and CXCL9). B. Transcription factor (TF) adaptation model. C. ODE model without adaptation (purple dotted lines) and with adaptation (orange dashed lines) best fit to IRF1, CXCL10, and CXCL9 population mean data (black solid lines) in all IFNγ dose and duration conditions. Prior to fitting, the fluorescence data here was rescaled to peak at the same level (about 40) for each gene.

While this deterministic model does a reasonably good job describing the mean response of the cells, it cannot capture the cell-to-cell variability that we see in our data. We were particularly interested in the striking heterogeneity in CXCL10 expression, especially given its relatively early expression following IFNγ stimulation. Our hypothesis was that this variability for CXCL10 may be due to slow and strongly stochastic chromatin dynamics. To test this hypothesis, we developed a stochastic model to model the dynamics of individual cells and characterize the distribution of responses. We used the same species and reactions as in our deterministic model but treated them stochastically using direct Gillespie algorithm (71) with the same rates as in our fitted deterministic model. While we did observe a strong variability in cellular CXCL10 time traces among individual runs in our simulations, we also saw that, given long enough stimulation time, all cells eventually express CXCL10 within the timeframe of our experiment (Fig. 5B). This results in a narrow distribution of maximal CXCL10 expression in the population of cells, contrary to what we see in experiments where even for continuous stimulation there is a significant fraction of cells that never express CXCL10 (Fig. 5E).

**Figure 5.**
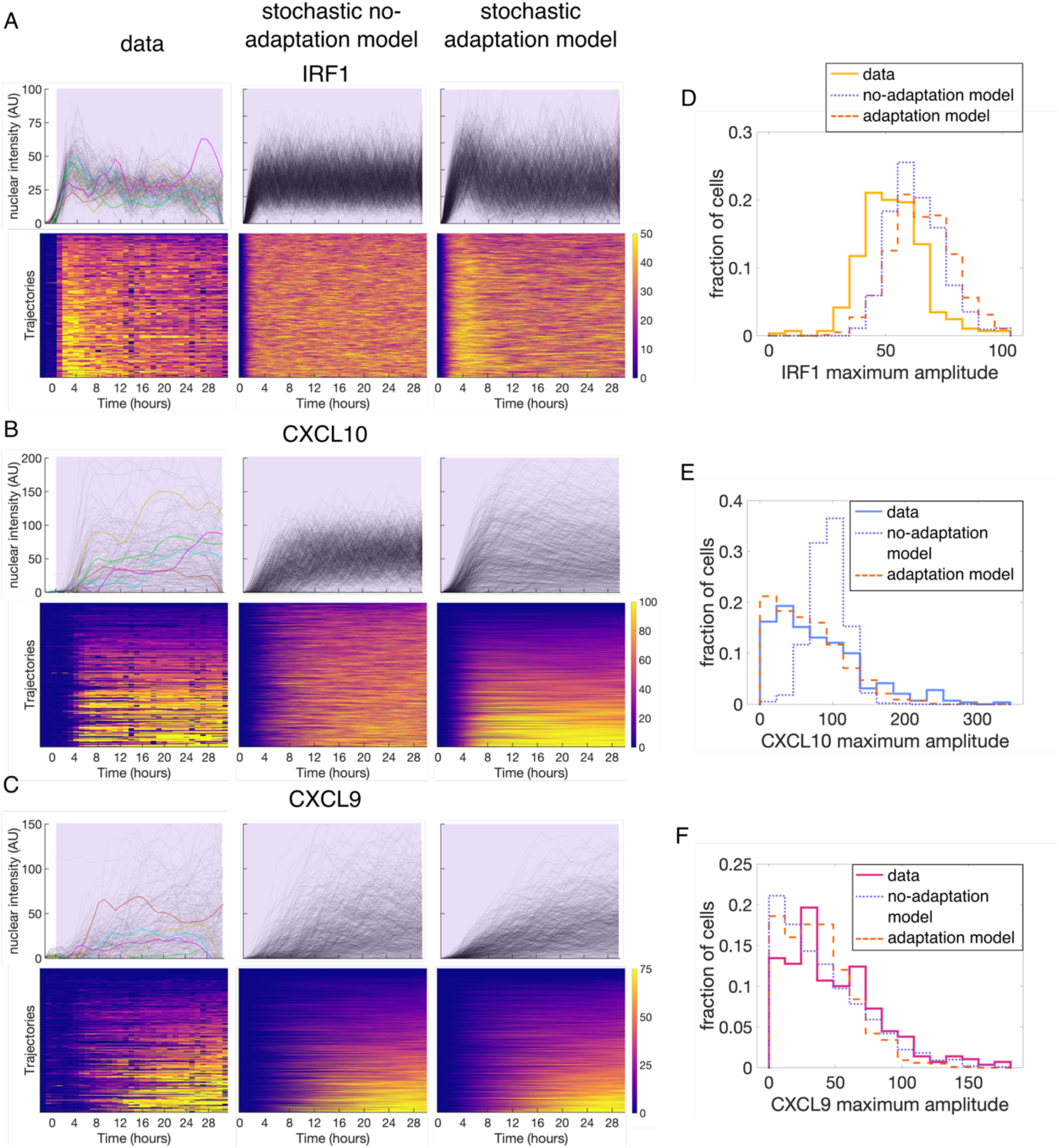
Stochastic modeling of IFNγ-induced gene expression. A-C. Comparison of experimental and stochastic simulation results for both no-adaptation and adaptation models for constant 10 ng/mL IFNγ stimulus for IRF1 (A), CXCL10 (B), and CXCL9 (C). The top panels show the fluorescence of each cell as a grey line and the bottom row panels show the same data in a heatmap. Heatmaps are sorted by maximum value for each gene individually. D-F. Histograms showing distribution of IRF1 (D), CXCL10 (E), and CXCL9 (F) maximum amplitude both experimentally (solid line) and in the no-adaptation (purple dotted line) and adaptation (orange dashed line) stochastic models for constant stimulus of 10 ng/mL IFNγ.

There could be two explanations for this discrepancy. One possible explanation is that there is a stable sub-population of non-responding cells that have CXCL10 permanently silenced at the chromatin level and thus never express CXCL10. The other possible explanation is that there is adaptation of the upstream signaling pathway, so after a certain time the stimulation of the downstream gene expression gets suppressed. Thus, even for a persistent upstream IFNγ stimulation, cells only have a finite time window in which they can express CXCL10 in response to this stimulation. Previous studies have shown that constant IFNγ stimulation leads to a peak of STAT1 nuclear translocation around 0.5-1 hour after stimulation, followed by STAT1 slowly leaving the nucleus, and that STAT1 nuclear localization is essential for its transcription factor activity (45, 72). We have also confirmed these STAT1 dynamics in our system using immunofluorescence for STAT1 (Fig. S4B). We have further experimental evidence for adaptation in our IRF1 data, where we see that upon constant IFNγ stimulation IRF1 levels decrease from their peak at a rate much slower than their putative degradation rate and in fact IRF1 remains expressed for as long as stimulus is present. This can best be explained by continued production of IRF1 at a substantial but sub-maximal rate due to upstream signal adaptation (Fig. 4C).

To account for this adaptation of the upstream signal, we modified our stochastic model to include a STAT1-induced negative regulator that inhibits STAT1in a negative feedback manner (Fig. 4B and S4A). This modification sets an upper limit on the time by which CXCL10 must be expressed in order to be expressed at all, and this results in the cell amplitude distribution looking more similar to the data (Fig. 5E). Adaptation also improves the stochastic model fit to the data for CXCL10 and CXCL9 lag time as well (Supp. Fig. S5B and C). We also incorporated this stimulus adaptation back into our deterministic model and observed that this greatly improved the agreement between the model and the data, especially for IRF1 under constant stimulation (Fig. 4C). When we use the stochastic model with adaptation to compute the distribution of maximum amplitude and lag time across all amplitude and duration conditions, we see that the histograms match the experimental data for most conditions (Fig. S5D). From this we conclude that a combination of the slow chromatin-opening step and the upper limit on the time for chromatin opening set by the timescale of STAT1 adaptation can produce the protein expression of varying amplitude but defined timing that we see from CXCL10, and that this can also produce the CXCL10 low-to-non-responders that we see even under constant stimulation.

### Response to repeated IFNγ pulses suggests slow, stochastic chromatin opening controls CXCL10 gene expression

While our model with adaptation describes the data for persistent and single-pulse stimulation quite well, the data reported thus far cannot eliminate the possibility that CXCL10 is simply epigenetically silenced in the low-to-non-responding cells. We also cannot rule out cell cycle effects, as previous studies have revealed that there are genes whose expression is biased towards specific cell cycles stages and thus cell cycle could be a source of gene expression variability (2, 73–77). To assess these possibilities, we performed further experiments and analyses. To evaluate the dependence of CXCL10 expression on cell cycle progression, we grouped cells by approximate cell cycle stage at the onset of IFNγ stimulation and observed that cells in all stages of the cell cycle respond similarly (Fig. S6A and B), thus ruling out the cell cycle as a driver of the observed heterogeneity.

To test whether CXCL10 was permanently epigenetically silenced in the low-to-non-responder cells, we decided to expose our cell populations to two pulses of IFNγ separated by a long enough interval so that any possible adaptation would be recovered. If the cells had permanently silenced chromatin, the cells that did not respond to the first pulse would also not respond to the second pulse. However, if the lack of response is due to the stochastic event of chromatin being closed during a pulse, then some cells that are silent during the first pulse could still respond during the second, and vice versa. We first tested this scenario in our stochastic model with adaptation, where we modeled stimulating the cells with two 4-hour pulses of IFNγ with a 10-hour off-interval. Our model predicted that all cells would show uniform expression of IRF1 in response to each input pulse, which is a consequence of relatively fast chromatin dynamics for IRF1 (Fig. 6A and B). However, for CXCL10, the chromatin dynamics are much slower, and so some of the cells that did not express CXCL10 in response to the first pulse still expressed CXCL10 in response to a second pulse (Fig. 6A and C). This CXCL10 result contrasts with what we would see if CXCL10 is stably silenced, where non-responding cells would remain non-responding to both pulses. The model also predicts low CXCL9 expression, as expected for such a short stimulus duration, but that the CXCL9 response to the second pulse will be slightly higher than that to the first pulse given that the chromatin at the CXCL9 locus closes slowly.

**Figure 6.**
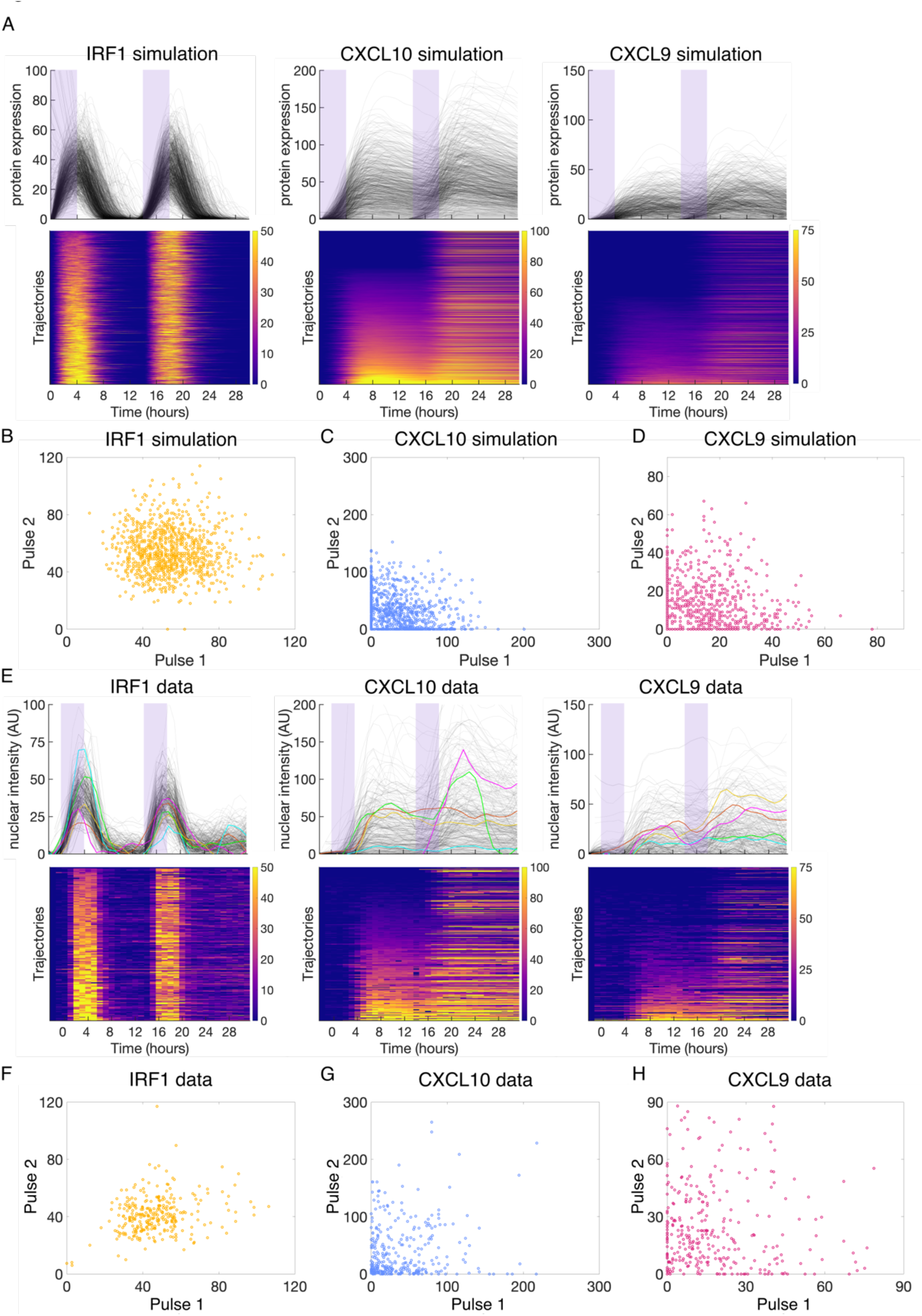
Model simulation and experimental results for cells exposed to two pulses of IFNã stimulation. A. Stochastic adaptation model results of 1000 cells exposed to two 4-hour pulses of 10 ng/mL IFNã with a 10-hour off period between the pulses. Line plots show each cell as a line and heatmaps show each cell as a row. Heatmaps are sorted by maximum florescence in the first 14 hours after the first onset of stimulation. B-D. The maximum protein expression amplitude change that each simulated cell in A achieved in each pulse window (from pulse onset to 10 h after the pulse ended) plotted against each other for IRF1 (B), CXCL10 (C), and CXCL9 (D). E. Experimental data of cells exposed to two 4-hour pulses of 10 ng/mL IFNã with a 10-hour off period between the pulses. Line plots show each cell as a line and heatmaps show each cell as a row. Heatmaps are sorted by maximum florescence in the first 14 hours after the first onset of stimulation. F-H. The maximum amplitude of expression that each experimental cell in C achieved in each pulse window plotted against each other for IRF1 (F), CXCL10 (G), and CXCL9 (H).

We performed the corresponding two–pulse experiment with the same protocol and observed single-cell responses that are quantitatively consistent with our model simulations (Fig. 6 E-H). These results rule out the possibility that CXCL10 is stably silenced in a subpopulation of cells and support our model, in which CXCL10 expression is controlled by slow, stochastic chromatin opening in response to an input signal that adapts around four hours, slightly longer than the timescale of chromatin opening. When we categorized the cells as responding to the first, second, both, or neither pulses, the probability of responding to each pulse is similar regardless of whether the cell responds to the other pulse, further supporting the stochastic nature of gene activation as predicted by our model (Supp. Table 3). We also see that the data match the model predictions for IRF1 and CXCL9, specifically that as predicted by the model the CXCL9 response to the second pulse is somewhat higher than to the first pulse (Fig. 6 B, D, F, H).

We further analyzed the quantitative features of the CXCL10 response to repeated input. To do this, we categorized cells as responding to the first (+/-), second (-/+), both (+/+), or neither (-/-) pulses. When we compare the response to the first pulse of stimulus between +/+ cells and +/-cells, we see that both the amplitude and slope of the response in +/- cells is higher than that of +/+ cells (Fig. S6C, D). When we compare the response to the first and second pulses in +/+ cells, we see that the slope of the second response is higher than that of the first response (Fig. S6E). Comparing the lag time, slope, and amplitude between the two pulses in single +/+ cells also shows that the majority of cells have a higher slope in their response to the second pulse (Fig. S6G). These results suggest that while CXCL10 activation is random, the expression level and slope can be affected by the cell’s expression history.

To further confirm the stochasticity in CXCL10 expression, we compared IRF1, CXCL10, and CXCL9 expression in sibling cells. For siblings that divide between 11 h and 0 h before the addition of IFNγ, there is weak positive correlation between sibling pairs for expression of each of IRF1, CXCL10, and CXCL9, and the correlation coefficient for CXCL10 is lower than that of IRF1 (Fig. S6, I-K). This lack of strong correlation between siblings, especially for CXCL10, is in accord with our repeated pulse experiment, supporting the intrinsic stochasticity in gene activation as described in our model.

## Discussion

In this work, we have used endogenous fluorescent reporters to determine how three different IFNγ-inducible genes can decode dynamic IFNγ stimulus in divergent ways. We found that macrophages express IRF1 strongly in response to low-concentration or transient IFNγ stimulus. In contrast, IFNγ stimulus must be higher in concentration or longer in duration for macrophages to express CXCL10 and CXCL9. Mechanistically, these different ways of decoding dynamic stimulus point to different mechanisms underlying expression of each gene. Previous work has shown that, in the absence of signal, chromatin at the IRF1 locus is ready for transcription, while the CXCL10 locus is slightly open but not yet transcriptionally active and the CXCL9 locus has repressive chromatin marks (78–80). These basal chromatin states align with the fast, homogenous IRF1 expression that we see, as well as the delayed expression of CXCL10 and the even more delayed expression CXCL9. While our modeling provides a possible explanation of these delays due to the speed of chromatin remodeling, further experimental work is needed to elucidate the mechanisms of gene expression at each locus. For example, we find that treatment with A485, a p300 inhibitor, delays CXCL10 expression, pointing to p300 recruitment as important for the timing of CXCL10 expression (Supplemental Fig. 7A).

We observe relatively uniform IRF1 expression among single cells, but remarkable cell-to-cell variability in CXCL10 and CXCL9 expression. The heterogeneity seen for CXCL10 is particularly notable because CXCL10 is expressed relatively early and at a high level, which are characteristics generally associated with lower heterogeneity. Further, the existence of a subpopulation of low-to-non-responding cells even upon a constant saturating-level input is intriguing and cannot simply be attributed to slow chromatin kinetics. In contrast, the homogeneity in IRF1 expression and the heterogeneity in CXCL9 expression are expected for a fast primary response gene and a very delayed secondary gene, respectively.

Our modeling suggests that the unique features of CXCL10 heterogeneity can be explained by slow, stochastic chromatin opening being on a similar timescale to that of upstream signal adaptation (∼4 h). As a result, cells have a defined and limited time window in which to turn on CXCL10, and only a fraction of them will be able to open their chromatin in that window and initiate gene expression. In this way, the interplay of time scales for chromatin activation and upstream signal adaptation can enhance the heterogeneity in gene expression even for relatively fast-responding genes. This idea is further supported by our subsequent modeling and experimental analyses, showing that CXCL10 expression can be activated stochastically at each onset of stimulus with a similar distribution of high- and low-responding cells.

Our modeling also shows that the adaptation of upstream STAT1 signal is crucial for capturing the dynamic and heterogeneous features of gene expression. Future work will clarify the mechanisms of this negative regulation and identifying the specific molecular factors involved for each gene. In our modeling we see that the cooperativity factor for production of the negative regulator’s mRNA is less than 1, which is unusual and could point to additional gene regulatory steps in the production of the negative regulator. It is also likely that there are additional general and gene-specific forms of negative regulation in this signaling pathway in addition to the common-to-all-genes adaptation that we model here, as negative regulation is a key feature of pro-inflammatory signaling. For example, it is known that there are specific mechanisms to negatively regulate CXCL10 expression (81–83). Including these may improve the model fitting even more. It is also important to note that while our data are consistent with this model of slow, stochastic chromatin opening, there are likely additional mechanistic elements involved in CXCL10 expression.

Both the dynamics and heterogeneity of gene expression observed in this study could have important physiological relevance for macrophage functions. The differences in gene expression response to dynamic stimulus that we found may underlie the thresholds at which macrophages perform different tasks in response to pro-inflammatory stimulus. It is somewhat surprising that an upstream transcription factor such as IRF1 saturates its expression at such a low amplitude and duration of stimulus, but there are likely further levels of regulation for IRF1 downstream genes. The fast decay of IRF1 upon removal of stimulus could facilitate fast transcriptional changes when the stimulus is gone. In contrast to IRF1, CXCL10 and CXCL9 act as high-pass filters where they are only expressed in response to higher amounts of stimulus.

This may permit the macrophage to ascertain that there is a sufficient quantity of IFNγ to warrant recruiting T cells and further amplifying the immune response, so that the macrophage does not amplify the immune response unnecessarily. In this way, the expression dynamics of different genes enable a time-based coordination of various functions performed by macrophages in response to changing environmental signals. Future work could connect these dynamics to diseases where IFNγ signaling is perturbed.

The heterogeneity in gene expression may also be functionally relevant for macrophages. We speculate that the wide distribution in gene expression may allow for more precise tuning of total population CXCL10 output. In addition, it may be more robust to perturbations than a system where there were stable populations of CXCL10-expressing and CXCL10-non-expressing cells, as in this system a single cell can switch between expressing CXCL10 and not expressing CXCL10. Future work could engineer populations of macrophages to either respond as our cells do or to stably express CXCL10 at a specific level and to investigate how this changes T cell recruitment and immune response progression *in vivo*. Heterogeneity in CXCL10 expression among genetically identical cells has been seen previously (21, 31), and there have been reports of heterogeneity in expression of other secreted cytokines, which we also see here with CXCL9 (84). It would be interesting to investigate if high heterogeneity is a common feature of secreted cytokines in the immune system.

Our work also provides an extension from the single-cell work done in macrophage response to primary infections to investigate how macrophages respond to cytokines that they experience in the middle of an immune response rather than at the beginning. As one point of comparison, we see that under conditions of constant IFNγ, IRF1 levels oscillate in both our experiments and our modeling. This is reminiscent of the NFkB oscillations that are observed in response to certain stimuli and suggests that more work should be done on oscillations of pro-inflammatory transcription factors to see if this is a more general phenomenon.

## Acknowledgements

We thank the Murre, Pasquinelli, and Hasty labs for use of their equipment, and the Reck-Peterson lab for the gift of the STAT1 antibody. We thank Dr. Terry Hwa and the entire Hwa lab for providing critical feedback periodically throughout the project. Cell sorting was done at the Sanford Consortium Human Embryonic Stem Cell Core with the help of their staff. We thank Dr. Phuc Nguyen and Sumedha Ravishankar for critical feedback on the manuscript. This work was supported by the NIH-sponsored Quantitative Integrative Biology Training Grant (T32GM127235) on which BN was a trainee, and NIH F31AI161903 (to BN), NIH R01GM111458 (to NH) and NIH R01 GM144595 (to NH and LST). AVN is supported by the Simons Foundation through the Principles of Microbial Ecosystems (PriME) collaboration (Grant no. 542387). JS is supported by a PhD Fellowship from Boehringer Ingelheim Fonds.

## Author contributions

B.N. designed the study with input from N.H. B.N. and J.S. designed the CRISPR knock-in strategy. B.N. performed all experiments and analyzed the data. B.N., A.V.N, and L.S.T. did the modeling. B.N. wrote the initial draft of the manuscript. All authors edited and contributed to the final version of the manuscript.

## Methods

### Cell culture

RAW 264.7 cells were ordered from ATCC (cat # TIB-71) and cultured in Dulbecco’s Modified Eagle Medium (DMEM) (Cytiva HyClone cat# SH30022FS) supplemented with 4500mg/L glucose, 4.0mM L-Glutamine, 10% fetal bovine serum, and 1% penicillin/streptomycin. Cells were cultured at 37C, 5% CO2, and 90% humidity.

### IFNγ and A485 treatment

Cells were treated with interferon gamma (Prospec Bio #CYT-358) at the concentrations stated in the text; if not stated them the concentration is 10 ng/mL, which corresponds to 100 U/mL. For A485 treatment, cells were treated with 10 μM of A485 (Tocris #6387) for two hours before addition of IFNγ.

### Cell line construction

We used CRISPR/Cas9 genome editing to knock in fluorescent proteins at the endogenous locus of genes to tag them. We designed guide RNAs (gRNAs) for CRISPR/Cas9 genome editing using an online CRISPR tool (http://crispor.tefor.net/). Three guides were ordered from Eurofins genomics and cloned into pSpCas9(BB)-2A-Puro (Addgene #48139) plasmids using a one-step restriction-ligation protocol (85). To test the cutting efficiency of the gRNAs, we transiently transfected the gRNA plasmids into NIH3T3 cells using Lipofectamine 2000 at a 1:3 Lipofectamine : DNA ratio. Transfected NIH3T3 cells were selected with puromycin (1 ug/mL) for 2 days and then grown out to a confluent 10-cm plate (2-3 days). We extracted genomic DNA from these cells, amplified the 1kb region around the cut site using PCR, and sent the PCR amplicon for Sanger sequencing. The sequencing results were then analyzed with the Synthego ICE analysis tool (https://ice.synthego.com/<u>#/</u>) and the gRNA with the best cutting efficiency was chosen.

We designed donor plasmids to have ∼1kb of homology on each side of fluorescent protein insertion site, and used primer tails to synonymously mutate either the PAM site or the gRNA recognition sequence so that the knock-in allele cannot be edited again. We used PCR to amplify these homology arms from the genomic DNA, and to amplify the fluorescent proteins from plasmids (SYFP2 from pSYFP2-C1 Addgene #22878, mCerulean and mCherry from existing plasmids in the lab). Donor plasmids were assembled in a pUC19 backbone using Gibson assembly and confirmed by Sanger sequencing. A flexible linker sequence (amino acid sequence GDGAGLIN) was used between IRF1 and the SYFP2 tag, and a T2A sequence was used between the protein and fluorescent tag for CXCL10 and CXCL9 (86). This T2A sequence acts as a translational skip site so that the endogenous chemokine can be secreted into the media and the fluorescent protein can be retained in the cell and so measured. We added an SV40-NLS tag to the fluorescent proteins driven by CXCL10 and CXCL9 so that we could measure fluorescence intensity in the nucleus. The nuclear marker consists of an NLS-iRFP (87) driven by a constitutive EF1alpha promoter (gift from Jan Soroczynski) inserted into the Tigre locus (86). This nuclear marker greatly facilitates cell segmentation and tracking. Plasmids were purified for transfection using a Macherey-Nagel NucleoBond Xtra Midi EF kit. A full list of plasmids can be found below.

We used the Neon transfection system to transfect the cells. 2.5 x 10>6 RAW 264.7 cells were used per electroporation, along with 15ug of gRNA-Cas9 plasmid and 15ug of donor plasmid in a 100uL Neon tip using R buffer. The electroporation setting was 1680V, 20ms, 1 pulse, after which the cells were plated into a one well of a 6-well dish of antibiotic-free DMEM. Three identical transfections were done per condition. The transfected cells were then grown to a confluent 10-cm plate (7-9 days depending on cell viability after transfection), induced with IFNγ, and sorted on an Aria Fusion cell sorter to select for cells expressing the newly knocked-in fluorescence protein. Cells were sorted into conditioned media with 20% FBS in 96-well plates with 1 cell per well and then 12 clones were grown up and screened. The cell line used in this paper was created by knocking in the nuclear marker, then tagging IRF1, CXCL10, and CXCL9 sequentially and growing up single clones after each knock-in.

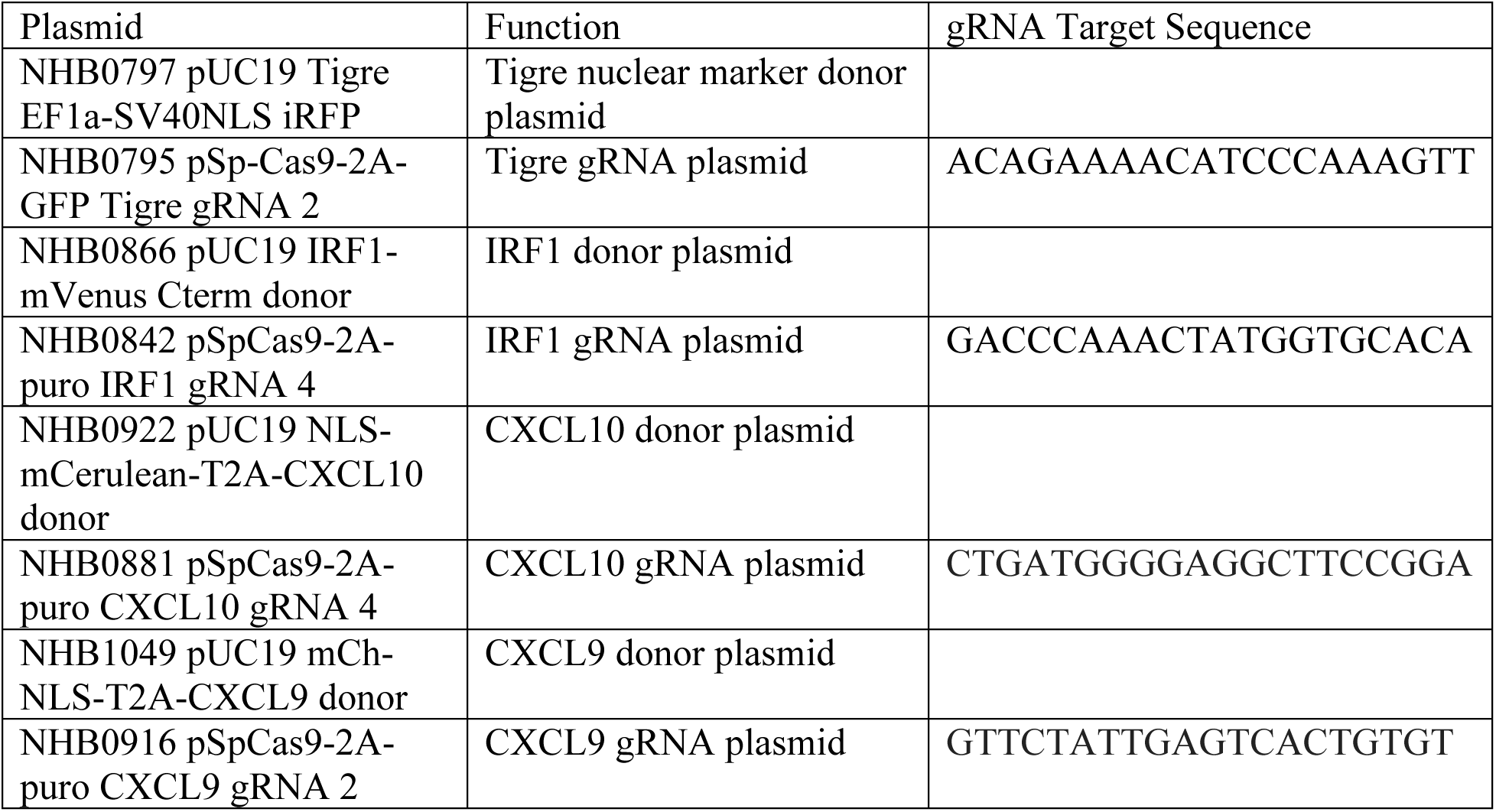

### Cell line screening

The 12 clones that were screened were first imaged on our microscope for fluorescence induction after IFNγ stimulation, with the goal being to choose a clone that was representative of the majority of the clones and had fluorescence matching what is known about the induction of that gene. The insertion region was also amplified by PCR and sequenced to identify any mutations that are present at that locus, and a clone was chosen that either had no mutation (CXCL10 and CXCL9, which are both homozygous tags), or else a mutation that did not affect protein function. The IRF1 tag is heterozygous. For IRF1, the tagged allele has perfect sequencing, and the non-tagged allele has a twelve base-pair deletion that makes it so what while the wild-type protein ends PSIQAIPCAP*, the untagged IRF1 in our cells ends PSIQAP*. However, this mutated allele produces the same level of IRF1 as a wild-type allele by Western blot and also induces IRF1 downstream genes to the same level as wild-type IRF1 by qPCR. There were no clones with IRF1 homozygously tagged, and this selected clone had the least affected wild-type allele. Cells from each clone were also induced with IFNγ and had qPCR done on the tagged gene and several other IFNγ-induced genes to make sure that the induction matched wild-type induction, and a clone was chosen where there was minimal difference in qPCR levels in the clone with tagged proteins vs the untagged wild-type cells.

### Microfluidic device fabrication

SYLGARD 184 silicone elastomer base was mixed with 10% of SYLGARD 184 silicone elastomer curing agent, degassed for 20 minutes, poured onto a custom wafer (68), degassed for 2 hours, and then baked at 80 C overnight. Individual PDMS chips were cut apart, had four holes punched in them, and cleaned with ethanol, water, and tape. Coverslips were cleaned with heptane, methanol, and water, and then dried using compressed air. Chips were bound to the coverslip in a UVO binder and then baked at 80 C overnight to bond permanently. Two individual chips were bound to each coverslip.

### Bulk plate-based assays

For bulk plate-based induction assays, used for qPCR and Western blots, a confluent 10 cm or 6 cm plate of RAW 264.7 cells was induced with the stated concentration of IFNγ. The cells were then collected after the induction time and pellets were frozen to be used in downstream qPCR or Western blots.

### qPCR

Total RNA was extracted from RAW 264.7 cells using Trizol. The RNA was then diluted to 100 ng/mL and converted to cDNA using a High Capacity cDNA Reverse Transcription Kit (applied biosystems, cat # 4368814). qPCR was then done using PowerUp SYBR Green master mix (Thermo Fisher cat # A25776) on a QuantStudio 3 thermocycler. Reactions were performed in triplicate and compared to uninduced RAW 264.7 cells to calculate the fold-change.

Primer sequences are as follows:

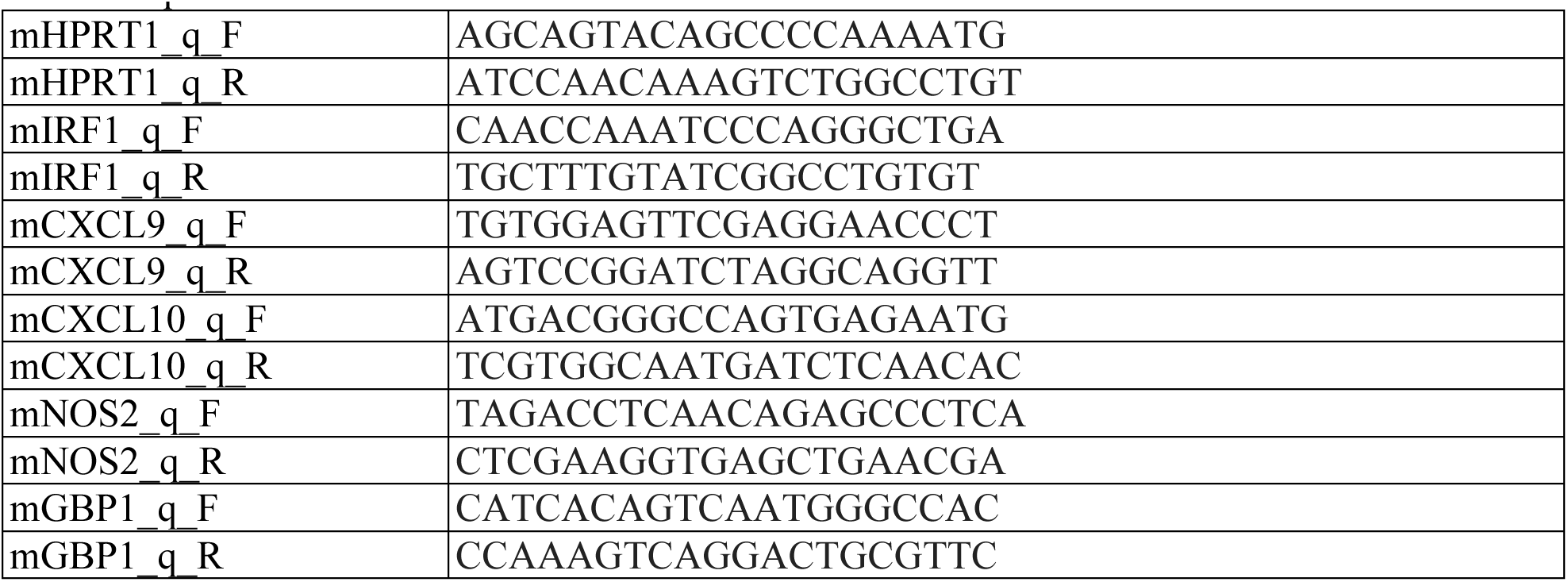

### Western blot

Proteins were extracted from RAW 264.7 cell pellets using IPH buffer sitting on ice for 30 mins. Western blots were run with a standard protocol. IRF1 antibody was D5E4 from Cell Signaling Technology.

### Immunofluorescence

RAW 264.7 cells were seeded onto coverslips in 24-well plates at a density of 4×10>4 cells per well. After 24 hours, the cells were induced with 10 ng/mL IFNγ at staggered times so that at the end of the experiment there were cells that had been induced for 24, 8, 6, 4, 2, 1, 0.5, and 0 hours. Cells were then washed with PBS, fixed in 4% paraformaldehyde for 15 minutes, and permeabilized with 0.5% Triton-X for 10 minutes. Cells were blocked for 30 minutes in 2% bovine serum albumin in 1X PBS, which was also used as the antibody diluent. STAT1 antibody D1K9Y (Cell Signaling Technology) was used at a concentration of 1:1000 and an incubation time of two hours, followed by secondary antibody incubation (Cell Signaling anti-rabbit IgG F(ab’)2 fragment Alexa Fluor 488 conjugate) for 1 hour. Slides were mounted in Prolong Diamond and cured overnight. Slides were imaged on a Nikon Eclipse Ti inverted microscope with a 500 ms exposure time for the far-red nuclear marker and a 300 ms exposure time on the GFP channel for STAT1.

### Imaging experiments

#### Plate-

Cells from a confluent 10-cm plate were seeded 20 hours before the experiment in 24-well plates at a density of 4×10>4 cells/well. Immediately before imaging, cells were washed with PBS and 500uL of phenol-red-free DMEM (Gibco cat# 31053028) with 4500mg/L glucose, 4.0mM L-Glutamine, 10% fetal bovine serum, and 1% penicillin/streptomycin was added to each well.

The plate was then brought to microscope and imaged for two hours before addition of stimulus, and then for 48 hours after. Stimulus was added with a micropipette while the plate remained on the microscope stage.

#### Microfluidic-

Each chip on the coverslip was vacuumed for 3 minutes before cell loading. A confluent 10-cm plate of RAW 264.7 cells was washed with PBS, collected in DMEM, centrifuged at 200g for 3 minutes, resuspended in 3 mL of phenol-red-free DMEM, and loaded into the pre-vacuumed microfluidic device using vacuum loading to a density of ∼20-30 cells/trap. Each chip with cells loaded was attached to a syringe with clear DMEM as well as a waste tubing and incubated in a standard tissue culture incubator at 37C, 5% CO2, with humidity for 20 hours between seeding and setting up on the microscope to allow cells to adhere to the coverslip. All syringes have a manual turn-valve controlling output from the syringe into the attached PTFE tubing (Cole-Parmer EW-06417-21), and this switching was controlled manually. Immediately before setting up on the microscope, a second syringe containing phenol-red-free DMEM with 10 ng/mL IFNγ was attached to the chip but closed so that the cells continue to be in a no-IFNγ environment.

The syringes and chip were then set up on the microscope, where the syringes sit outside the environmental chamber at 12 cm above the chip, and the waste lines also sit outside the chamber at 30 cm below the chip. The chip is in an environmental chamber controlling the temperature at 37 C, 5% CO2, and humidity. The image ROI was set at 300×300 pixels, which corresponds to the size of one trap so that every image contains only one trap. 13 traps for each chip were chosen for imaging and were chosen to have a good number of non-clumped cells and to not image two adjacent traps (this minimizes phototoxicity). Images were taken for at least two hours before onset of stimulation. At the onset of stimulation, the valve on the DMEM + IFNγ syringe was opened and the valve on the DMEM-only syringe was closed, and at the end of stimulation, the DMEM-only syringe was opened and the DMEM+IFNγ syringe was closed.

Waste was collected in a 50-mL conical tube and measured to confirm that there was proper flow through the device during the experiment, which is about 0.5 mL/hour. Experiments without the correct amount of flow were discarded.

#### All-

Cells were imaged on a Nikon Eclipse Ti inverted microscope at 37 C, 5% CO2, with humidity. Phase and iRFP nuclear marker images were taken every 10 minutes, while fluorescence images were taken every 60 minutes to minimize phototoxicity. Images were taken with a 20X/0.45 NA objective using the perfect focus setting, and a Photometrics Evolve 512Delta EMCCD camera using Nikon Elements software. Exposure times were 500 ms for iRFP, 500 ms for mCherry, 300 ms for SYFP, and 100 ms for mCerulean.

### Image processing

Images were exported from the Nikon Elements software as multi-page tifs and imported into CellProfiler. A CellProfiler pipeline was used to subtract the background using a rolling-ball algorithm. Nuclei were then identified from the iRFP nuclear marker images using a minimum cross entropy algorithm for the microfluidic chip and Otsu algorithm for the 24-well plate, and a mask was made from these segmentations. The images taken from plate experiments and microfluidic experiments differ in their level of background and sharpness of the edges of the nuclei, which was why different segmentation algorithms worked best for each type of experiment and why we chose to use two different algorithms. Except in fitting our ODE model, the plate and microfluidic data are not compared against each other, and in the ODE modeling there is an introduction of a scaling factor (see modeling section) to make the plate and microfluidic values comparable. Fluorescence was quantified in each channel for each nucleus, and composite images were created with the object numbers overlaid on each fluorescence channel for manual confirmation of fluorescence and tracking. The images and a spreadsheet with the data were exported. Custom Matlab scripts were then used to organize the images into folders. We used u-track (88) implemented in Matlab to track the nuclei, and the tracking data were then converted into a format where each trace consists of the mean fluorescence intensity value for that cell at each timepoint. We then took only traces that are complete over the first 200 images of the experiment (∼33 hours, so ∼31 hours after IFNγ addition) and analyzed these further. All cells analyzed have strong trackable nuclear marker signal, indicating that they are alive for the duration of the experiment.

### Quantification and statistical analysis Replicates

For dose experiments, all doses were run in the same plate, and these plates were run in triplicate. One representative plate is used for all analysis shown. Each condition for plate experiments has 578-1016 cells. For microfluidic duration experiments, all conditions were run at least five times, with one chip on the coverslip running the experimental condition and the other chip receiving four hours of stimulus as a control between days and chips. For each duration modulation condition, we combined all cells from 2-3 different runs of that condition and used that as the data for that condition. This results in each condition including 459-921 cells. The statistics on these combined datasets match the statistics for each individual run that makes up the dataset. These runs were chosen since they are representative of all runs and all had their paired 4-hour control looking similar. For the microfluidic two-pulse experiment, this experiment was run three times and a representative run was chosen to show in these figures.

The representative run chosen includes 367 cells.

### General feature extraction

Features were extracted from all cells that had trajectories complete over the entire 31 hours of the experiment. Maximum amplitude for all genes was calculated as the maximum amplitude over the first 24 hours after IFNγ addition minus the mean of the two first images before IFNγ was added (which we suppose to be noise/basal gene expression and thus unaffected by IFNγ). Lag time was calculated as the first time at which the fluorescence value was 1.5x higher than that cell’s baseline level (defined as the mean of the first two images before IFNγ was added as well as the image right when IFNγ was added)^1^. In all cells the values of these first three images (the two before IFNγ addition and the one right when IFNγ was added) were very similar. Width for IRF1 was defined as the width at half of the maximum amplitude defined as the difference between the maximum value and the baseline expression. As CXCL10 and CXCL9 are transcriptional reporters and therefore do not represent endogenous decay rates, we do not calculate width for them.

Histogram boxplots were created using distributionPlot (Jonas (2017). Violin Plots for plotting multiple distributions (distributionPlot.m) (https://www.mathworks.com/matlabcentral/fileexchange/23661-violin-plots-for-plotting-multiple-distributions-distributionplot-m), MATLAB Central File Exchange. Retrieved April 28, 2023) with bins edges held constant for all conditions in a plot, but each histogram scaled individually, so the shape of the histograms is comparable but individual bin height is not comparable. This was chosen because having individual bin height be comparable led to overlapping histograms between conditions as in some cases there was one condition with the majority of its density in the smallest bin.

For the 2-pulse experiment, maximum amplitude was taken to be the maximum amplitude from the time of IFNγ onset to 13 hours after IFNγ onset (4 hours of IFNγ + 9 hours of time off) minus the expression at the time of IFNγ onset. This was chosen so that each pulse would be measured independently for itself and previous pulses would not contribute to later pulses.

CXCL10 feature extraction

The slope and pulse amplitude (used only in two-pulse experiment) for CXCL10 are extracted by fitting the initial CXCL10 rise (defined as the first 13 hours after IFNγ addition) to *y* = (tanh(*a* ⋅ *x* + *B*)) ⋅ *c* + *d* and defining the slope as *a* ⋅ *c* and the pulse amplitude as 2 ⋅ *c*.

### Sibling cells

Siblings are manually verified that they are in fact siblings, and their division time is defined as the first time when the cells are two separate cells. The sibling cells used in this analysis received 10 ng/mL IFNγ stimulus in a 24-well plate.

### Cell cycle

Cells were imaged for 11 hours prior to addition of IFNγ (addition of IFNγ being defined as time 0), and only cells that divided at some point during the experiment either before or after IFNγ addition were included in this analysis to define their cell cycle stage. Cells were placed into bins based on when they divided, with bin 1 being hours -11 to -7, bin 2 being hours -6 to -1, bin 3 being hours 0-4, bin 4 being hours 5-9, bin 5 being hours 10-16, and bin 6 being hours 17-22. This corresponds roughly to bin 1 being in G1/S when IFNγ was added, bin 2 being in G1, bin 3 being in G2, bin 4 being in S, bin 5 being in G1, and bin 6 being in G2. From observing cells that divided twice during the experiment we see that cell cycle length for most cells is about 17 h, but in some cells can be as long as 26 h.

### Calculation of coefficient of variation

The coefficient of variation (CV) is defined as the standard deviation of a distribution divided by its mean. We calculate general CV (in the main figures) by taking the CV of the expression level of all cells at each timepoint and plotting this result over time for all conditions. Calculation of CV for individual features was done by taking the standard deviation of that feature in all cells divided by its mean.

### Modeling

The block diagram of our non-adaptive model is shown in Fig. 4A. In addition to the transcription factor (TF), for each gene of interest, it includes five species that are involved in eight reactions: mRNA (M), protein (P), and three possible states of the chromatin at the locus of each gene of interest: closed (C), open-uninitiated (OU) and open-initiated (OI). The transition rates among the chromatin states and the rates of mRNA and protein synthesis and degradation are shown in Fig. 4A. We assume that mRNA synthesis can occur only when chromatin is in the open/initialized state. For simplicity, we also assume that the chromatin cannot close when it is in the initialized state. The concentration of the transcription factor is a nonlinear function of the stimulus concentration [IFNγ],

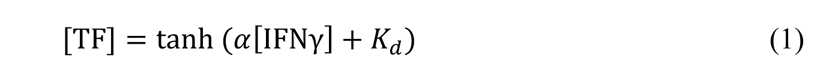

Chromatin opening rate and initiation rate are both proportional to [TF], while closing (*k*_2_) and deinitiation (*k*_4_) rates are constant.

In the generalized model that includes stimulus adaptation, we added two more species, regulatory mRNA (*regM*), and regulatory protein (*regP*), and 5 more reactions for the transcriptional negative feedback loop that suppresses the transcription factor (TF), see Fig. 4B.

For deterministic modeling of the system without adaptation, we introduce mRNA and protein concentrations ([M] and [P], respectively) and the fraction of chromatin loci to be in each of the three states ([C], [OU], and [OI], respectively). The set of the ODEs for these variables reads as follows:

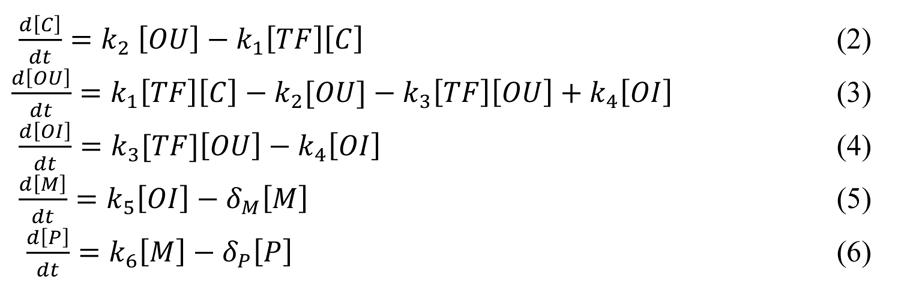

Because there are two chromatin loci and so there is a conservation [C]+[OU]+[OI]=2, one of these differential equations can be eliminated.

The deterministic model with adaptation involves, in addition to the five equations above, the following two differential equations

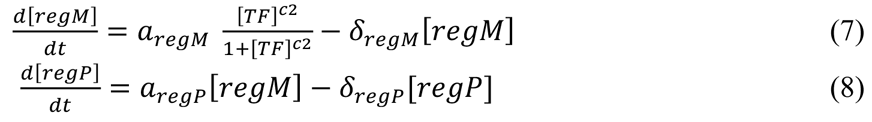

and the algebraic equation for the TF concentration that replaces (1):

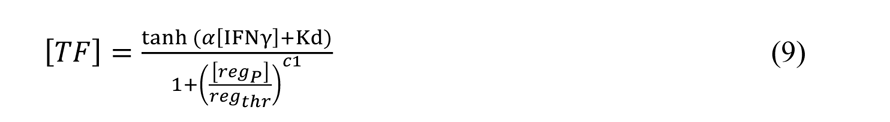

We used these deterministic models for all three genes of interest (IRF1, CXCL9, and CXCL10). To fit the model, we used the data for the mean values of the fluorescence for each gene at each time in each dose and duration condition. Since the conversion factor between the concentrations of proteins and the measured levels of fluorescence can differ between fluorescent proteins, before fitting we rescaled the data such that the mean expression peaks at 40. Since the amplitude of fluorescence in the plate experiments (used for the concentration experiments) is higher than in the microfluidic device (used for duration experiments), we scaled the microfluidic data based on the fact that the 10 ng/mL IFNγ plate condition should be equivalent to the 10 ng/mL constant IFNγ microfluidic condition. We calculated the scaling factor for each gene by dividing the maximum fluorescence in the 10 ng/mL IFNγ plate condition by the maximum fluorescence in the 10 ng/mL IFNγ constant microfluidic condition, and then multiplied all microfluidic data values for that gene by this scaling factor. The scaling factor for IRF1 was 1.7757, for CXCL10 was 3.1105, and for CXCL9 was 3.3009. We used the MATLAB function `minsearchbnd` for constrained optimization (John D’Errico (2012). fminsearchbnd, fminsearchcon (https://www.mathworks.com/matlabcentral/fileexchange/8277-fminsearchbnd-fminsearchcon), MATLAB Central File Exchange. Retrieved April 13, 2023) to fit the models for each of the three proteins of interest (IRF1, CXCL10, CXCL9). The best fit parameters for each of these cases are given in Supplemental Tables 1 and 2.

We note that there is a partial indeterminacy in the parameters obtained from fitting the deterministic model^2^ and parameters in Supplemental Tables 1 and 2 were chosen to be in the range of plausible values, and such that the stochastic simulations roughly matched the observed variability.

For the stochastic simulations, we used the same species and reactions as in the deterministic models. We employed the direct Gillespie algorithm (71) to compute the numbers of molecules of each species as a function of time. For the reaction rates we used the same parameter values that were obtained by fitting deterministic models (Supplemental Tables 1 and 2). After generating multiple stochastic runs (∼1000 per each experimental condition), we performed statistical analysis of simulation data by computing the mean values, standard deviations, and the distributions of selected features of the runs (maximum amplitude, lag time, etc.).

## Footnotes

1 For CXCL10, we can also extract lag time from these curves by fitting them to y = tanh(a*b)*c+d and defining lag time as – (1 + *b*)/*a*, which is the time when a line with slope *a* ⋅ *c* through the inflection point intersects with the baseline expression level. However, here do not use this calculation since it does not apply to IRF1 and CXCL9. Comparing these two methods of calculating lag time for CXCL10 provides very similar results. For the two-pulse experiment feature extraction we consider responding cells to be cells that fit this tanh function well, and non-responding cells to fit it poorly.

2 Given only a single species (fluorescent protein) for fitting, the dynamics for this species can be invariant with respect to suitable rescaling of pairs of reaction parameters, for example, by increasing translation rate and simultaneously decreasing transcription rate by the same factor. Using constraints on the expected rates of synthesis and degradation of mRNA and proteins we were able to narrow down this degeneracy, but to completely eliminate it, more detailed multivariate data would be necessary.

