## Supplemental Figures for "Quantifying dynamic pro-inflammatory gene expression and heterogeneity in single macrophage cells"

Supplemental Figure 1

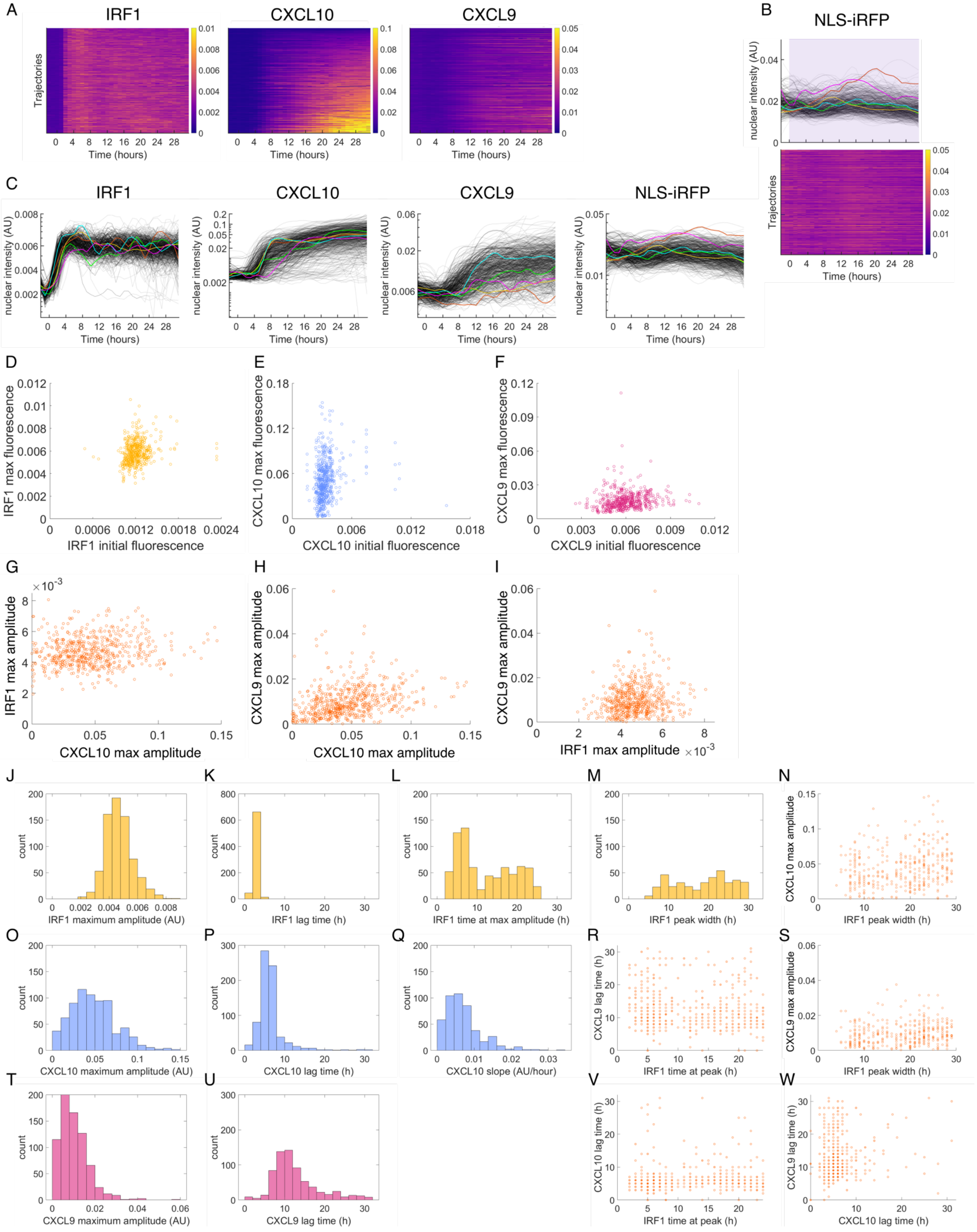

**Supplemental Figure 1**

A. Heatmaps of single-cell IRF1, CXCL10, and CXCL9 expression responses to 10 ng/mL IFNγ in a 24-well plate sorted in the same order for all three genes. B. Nuclear marker NLS-iRFP expression in response to 10 ng/mL IFNγ stimulus in a 24-well plate, shown both as single-cell traces and as a heatmap with each row being one cell. C. IRF1, CXCL10, and CXCL9 gene expression responses to 10 ng/mL IFNγ in a 24-well plate, plotted on a log scale. Each grey line is a cell, with five exemplary traces in color. The data visualized is the same as Figure 1D. D. Scatterplots correlating the maximum amplitude of gene expression over the first 24 hours of IFNγ stimulation (the raw maximum amplitude, not the maximum amplitude with the baseline subtracted off that we use later for downstream analysis) with the initial fluorescence before IFNγ was added. G-I. Scatterplots correlating the maximum amplitude of the gene expression over the first 24h post-stimulus between different genes in the same cell. Each point represents one cell. J-M. Histograms of IRF1 features for cells exposed to 10 ng/mL IFNγ for 31 hours in a 24-well plate. O-Q. Histograms of CXCL10 features for cells exposed to 10 ng/mL IFNγ for 31 hours in a 24-well plate. T-U. Histograms of CXCL9 features for cells exposed to 10 ng/mL IFNγ for 31 hours in a 24-well plate. N, R, S, V, W. Correlation of features of different genes in the same cells. For N, S each point represents one cell. For R, V, W each point represents one cell, and the opacity of the circle fill represents the number of cells at the same vertex.

Supplemental Figure 2

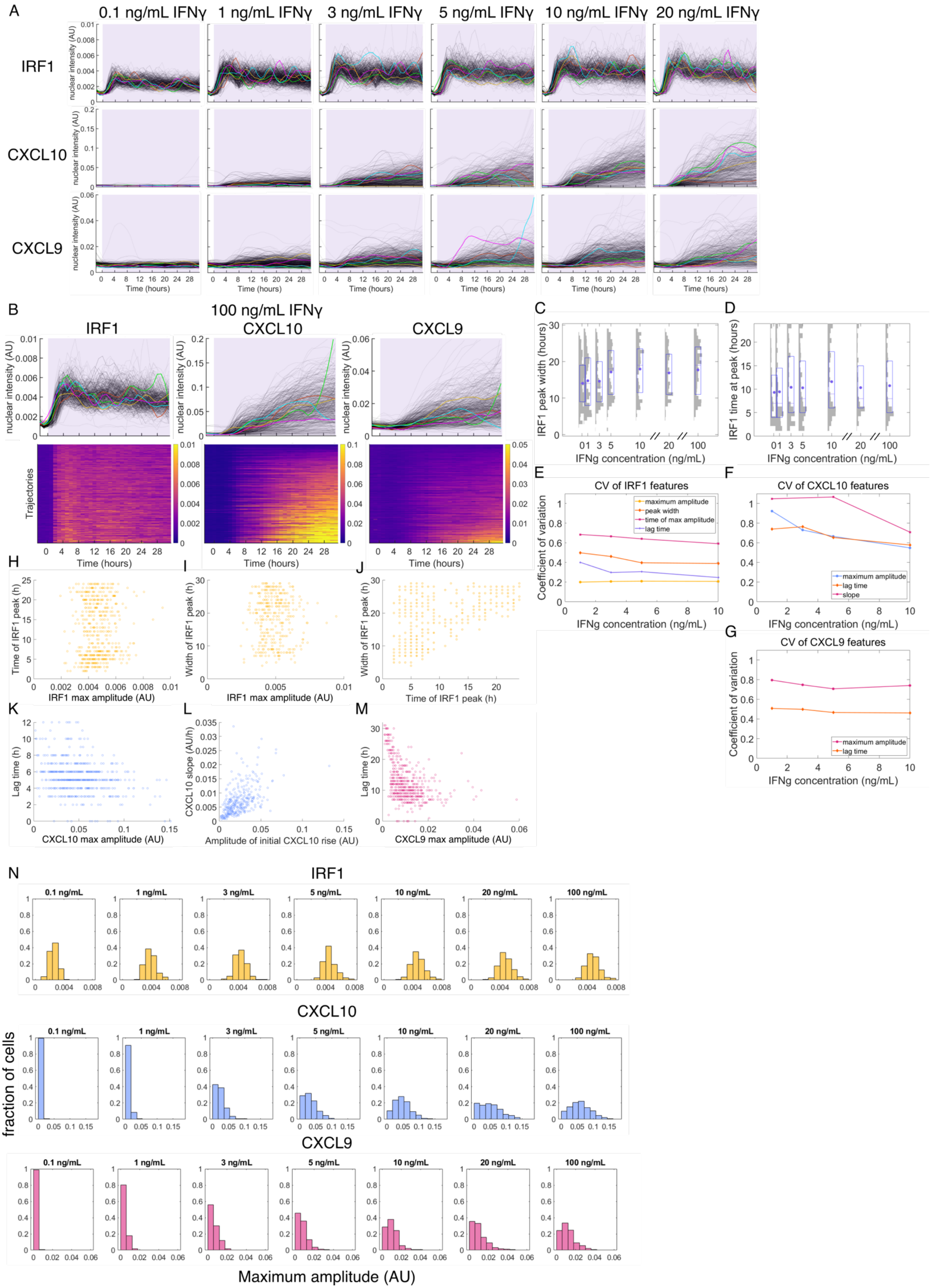

**Supplemental Figure 2**

A. IRF1, CXCL10, and CXCL9 gene expression responses to 1-20 ng/mL IFNγ in a 24-well plate. Same source data as Figure 2A-C but shown as single-cell traces. Each grey line is a cell, with 5 traces highlighted as examples. B. IRF1, CXCL10, and CXCL9 gene expression responses to 100 ng/mL IFNγ in a 24-well plate. In the top row, each grey line is one cell, with five exemplary traces highlighted. Bottom row shows the same data as heatmaps with each row representing one cell, and each heatmap sorted by single-cell maximum value for that specific gene. Purple shading indicates when the cells are exposed to IFNγ. C-D. Histograms overlaid with box plots showing the IRF1 peak width (C) and IRF1 time at peak (D) in single cells for each IFNγ concentration. Purple dot is mean, middle purple line is median, and purple box is the $25^{th}$-$75^{th}$ percentile. Grey shading shows the histogram distribution among single cells for each condition. E-G. Coefficient of variation for additional response features for IRF1 (E) CXCL10 (F) and CXCL9 (G). H-J. Scatterplots correlating features (H – amplitude vs peak time, I – amplitude vs peak width, J – peak time vs peak width) of IRF1 expression in response to 10 ng/mL IFNγ stimulus in a 24-well plate. K-L. Scatterplots correlating features (K – amplitude vs lag time, L – amplitude vs slope) of CXCL10 expression in response to 10 ng/mL IFNγ stimulus in a 24-well plate. M. Scatterplot correlating maximum fluorescence and lag time of the CXCL9 response to 10 ng/mL IFNγ stimulus in a 24-well plate. In the scatterplots, each dot is one cell and the opacity of the circle represents the number of counts. N. Histograms showing the distribution of maximal expression amplitudes in single cells in response to 0.1-100 ng/mL IFNγ.

Supplemental Figure 3

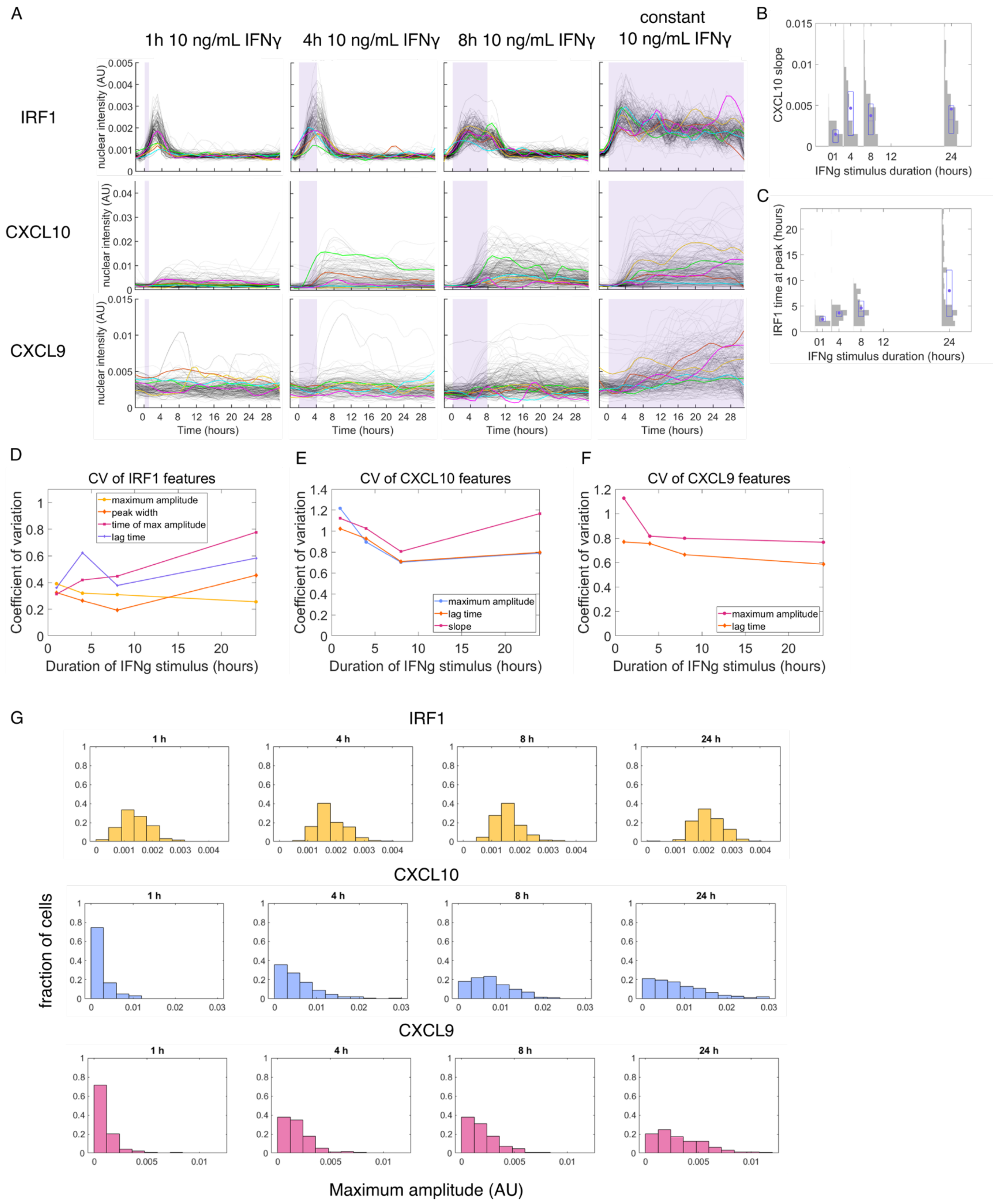

**Supplemental Figure 3**

A. IRF1, CXCL10, and CXCL9 gene expression responses to 1 h, 4 h, 8 h, and constant 10 ng/mL IFNγ stimulation in a microfluidic device. Same source data as Figure 3 A-C but shown as single-cell traces. Each grey line is a cell, with 5 traces highlighted as examples. B-C. Histograms overlaid with box plots showing the CXCL10 slope (B) and IRF1 time at peak (C) in single cells for each IFNγ duration. Purple dot is mean, middle purple line is median, and purple box is the $25^{th}$-$75^{th}$ percentile. Grey shading shows the histogram distribution among single cells for each condition. D-F. Coefficient of variation for additional response features for IRF1 (D), CXCL10 (E) and CXCL9 (F). G. Histograms showing the distribution of maximal expression amplitudes in single cells in response to 1 h, 4 h, 8 h, and constant 10 ng/mL IFNγ in a microfluidic device.

Supplemental Figure 4

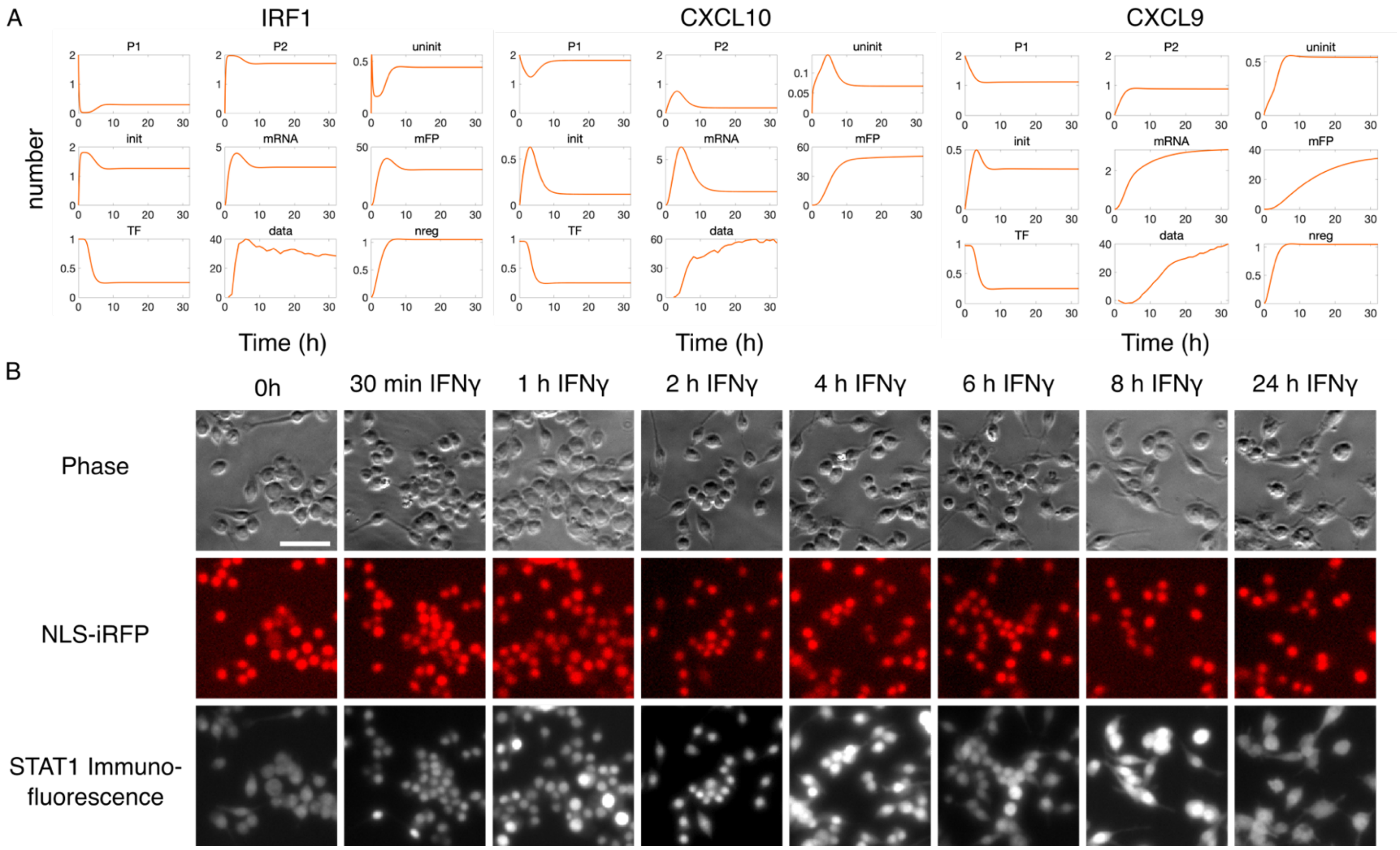


**Supplemental Figure 4**

A. ODE adaptation model intermediates for each of IRF1, CXCL10, and CXCL9 in the 10 ng/mL constant IFNγ stimulus condition. B. Immunofluorescence for STAT1 in RAW264.7 cells treated with 0-24 h of IFNγ. Scale bar on top left image represents 50 μm.

Supplemental Figure 5

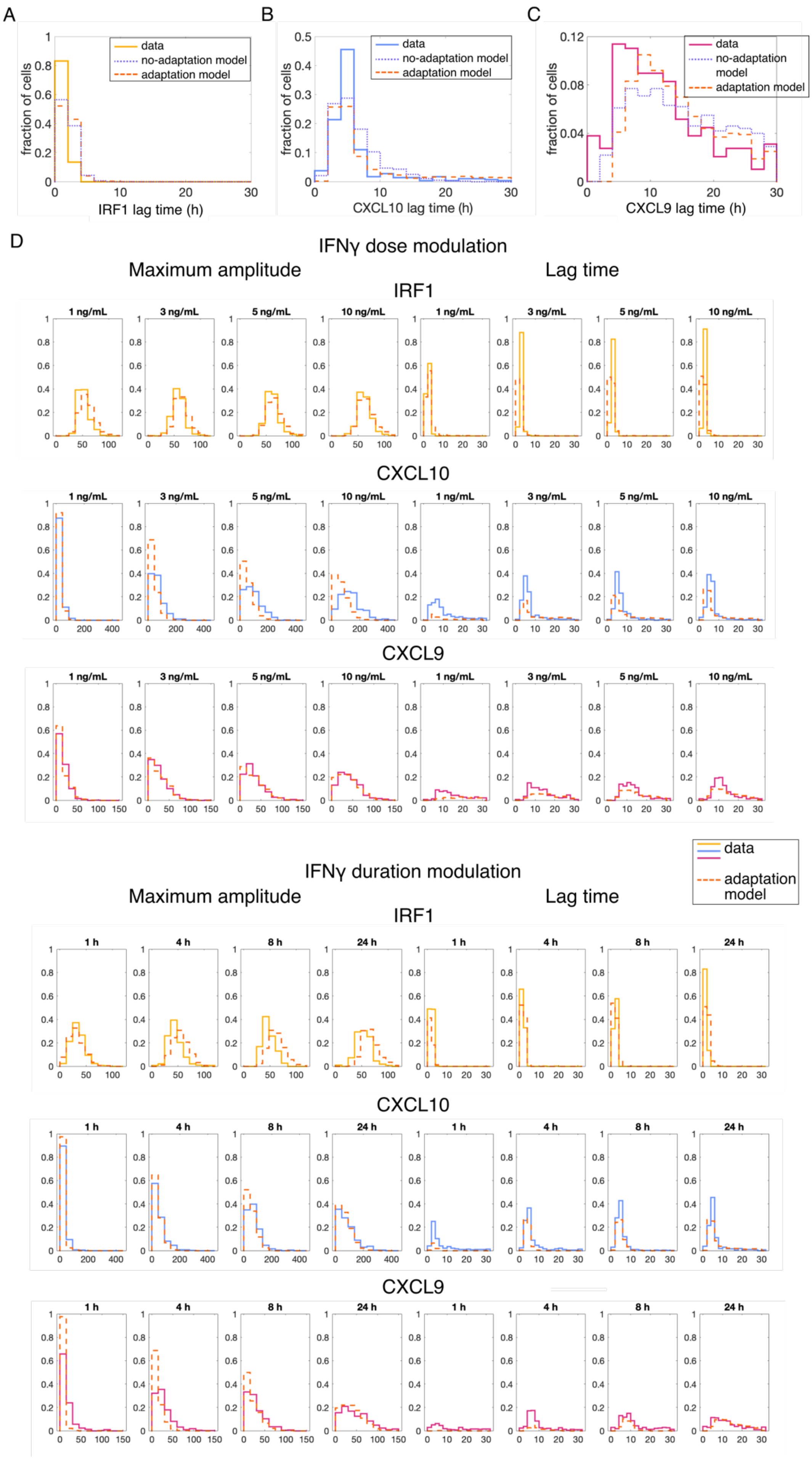


**Supplemental Figure 5**

A-C. Histograms showing distribution of IRF1 (A), CXCL10 (B), and CXCL9 (C) lag time both experimentally and in the no-adaptation (purple dotted line) and adaptation (orange dashed line) stochastic models for constant stimulus of 10 ng/mL IFNγ. D. Histograms showing distribution of maximum amplitude and lag time for IRF1, CXCL10, and CXCL9 for various dose and duration conditions in both the data (solid outline) and adaptation stochastic model (dashed orange line).

Supplemental Figure 6

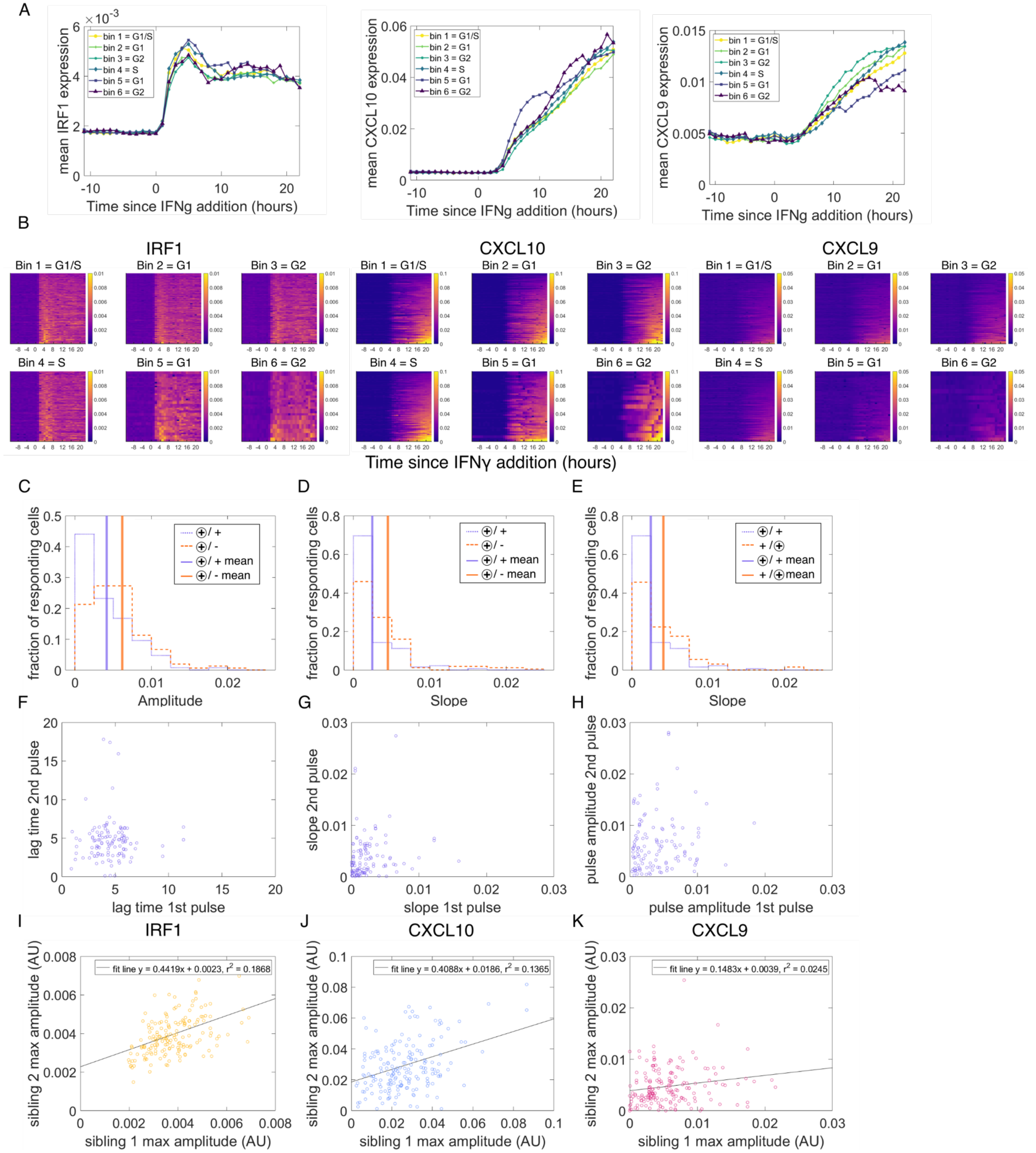

**Supplemental Figure 6**

A. Mean expression of IRF1, CXCL10, and CXCL9 in each cell cycle bin. B. Heatmaps showing expression of IRF1, CXCL10, and CXCL9 in each single cell, separated by cell cycle bin. Each row is one cell, and each heatmap is individually sorted by maximum expression. C and D. Histogram showing distribution of CXCL10 expression amplitude (C) and slope (D) in response to the first pulse of IFNγ between cells that respond to both pulses (+/+) and cells that only respond to the first pulse (+/-). The circled symbol shows the pulse that is being plotted. Solid lines show means of each distribution. Both the means (via t-test) and distributions (via KS test) are statistically significantly different at $p<.01$. E. Histogram showing distribution of CXCL10 expression slope in response to both the first and second pulse of IFNγ in cells that respond to both pulses (+/+). Solid lines show means of each distribution. Both the means (via t-test) and distributions (via KS test) are statistically significantly different at $p<.01$. F-H. Scatterplot of single cells that respond to both pulses (+/+) comparing the CXCL10 lag time (F), slope (G), and amplitude (H) in response to each pulse. I-K. Scatterplot showing the maximum IRF1 (I), CXCL10 (J), or CXCL9 (K) amplitude in the first 13 hours post-stimulus between sibling cells that divided in the 11 hours prior to stimulus onset.

Supplemental Figure 7

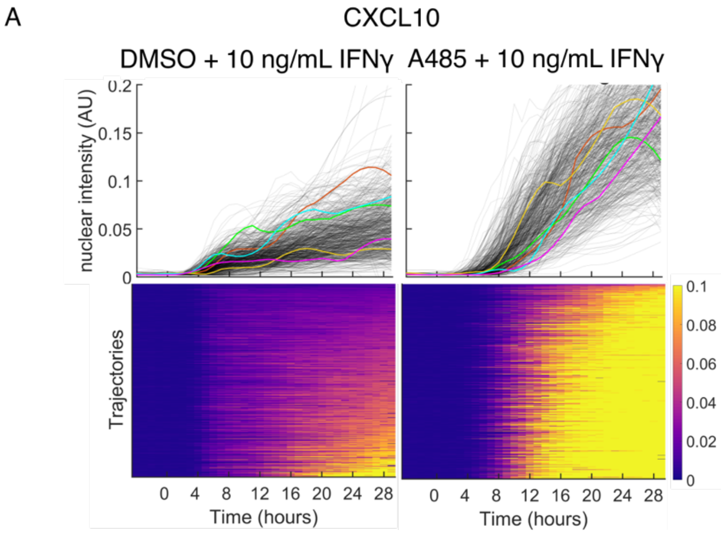


**Supplemental Figure 7**

A. CXCL10 expression in response to two hours of treatment with either DMSO or 10 µM of A485 followed by addition of 10 ng/mL IFNγ. Top row shows single-cell traces, each grey line is a cell, 5 traces are highlighted as examples. Bottom row shows the same data as heatmaps with each row being a cell.

**Supplemental Table 1. Parameters from fitting the deterministic ODE model both with and without adaptation to the data for each gene**

| | IRF1 no adaptation | IRF1 adaptation | CXCL10 no adaptation | CXCL10 adaptation | CXCL9 no adaptation | CXCL9 adaptation |
|---|---|---|---|---|---|---|
| $k_1$ | 10 | 6.2065 | .9188 | .3000 | .1217 | .18 |
| $k_2$ | 3.0923 | 1.0454 | 2 | 2 | 1.5894 | .0927 |
| $k_3$ | 4.0999 | 10 | .7269 | 8.1355 | 2.0036 | 10 |
| $k_4$ | .5884 | .8913 | .0435 | 1.1367 | 3.9950 | 4.0301 |
| $k_5$ | 4.7614 | 2.9586 | 2.7842 | 7.7541 | 9.2615 | 1.2306 |
| $k_6$ | 14.7298 | 10.7327 | 9.0105 | 1.3862 | 1.2439 | 1.5568 |
| $\delta_M$ | 1.9363 | 1.1467 | .8501 | .6265 | .1055 | .1329 |
| $\delta_P$ | 1.9363 | 1.1467 | .8501 | .0393 | .1716 | .1330 |
| $K_d$ | .0434 | .0359 | .025 | .025 | .1813 | .0873 |
| $\alpha$ | 3.2926 | 4.2873 | 1.5 | 1.9256 | 1.9865 | 1.9710 |

**Supplemental Table 2. Parameters used for adaptation of input IFNγ signal in deterministic ODE model**

| | |
|---|---|
| $a_{regM}$ | 3.8746 |
| $\delta_{regM}$ | .6531 |
| $a_{regP}$ | .2710 |
| $\delta_{regP}$ | .6530 |
| $reg_{thr}$ | .8792 |
| *c1* | 6 |
| *c2* | .2144 |

**Supplemental Table 3. Fraction of cells that do or do not express CXCL10 and CXCL9 in response to two four-hour pulses of IFNγ**

By CXCL10 fitting:

| | | |
|---|---|---|
| Fraction of cells that respond to the first pulse | Out of all cells | .7493 |
| | Out of cells that do respond to the second pulse | .6944 |
| | Out of cells that don't respond to the second pulse | .8021 |
| Fraction of cells that respond to the second pulse | Out of all cells | .4905 |
| | Out of cells that do respond to the first pulse | .4545 |
| | Out of cells that don't respond to the first pulse | .5978 |
| Fraction of cells that respond to both pulses | Out of all cells | .3406 |
| Fraction of all that respond to first * fraction of all that respond to second | | .3675 |

By CXCL10 threshold:

| | | |
|---|---|---|
| Fraction of cells that respond to the first pulse | Out of all cells | .7629 |
| | Out of cells that do respond to the second pulse | .6615 |
| | Out of cells that don't respond to the second pulse | .8743 |
| Fraction of cells that respond to the second pulse | Out of all cells | .5232 |
| | Out of cells that do respond to the first pulse | .4536 |
| | Out of cells that don't respond to the first pulse | .7471 |
| Fraction of cells that respond to both pulses | Out of all cells | .3406 |
| Fraction of all that respond to first * fraction of all that respond to second | | .3991 |

By CXCL9 threshold:

| | | |
|---|---|---|
| Fraction of cells that respond to the first pulse | Out of all cells | .5493 |
| | Out of cells that do respond to the second pulse | .3400 |
| | Out of cells that don't respond to the second pulse | .5973 |
| Fraction of cells that respond to the second pulse | Out of all cells | .1862 |
| | Out of cells that do respond to the first pulse | .1214 |
| | Out of cells that don't respond to the first pulse | .2727 |
| Fraction of cells that respond to both pulses | Out of all cells | .0633 |
| Fraction of all that respond to first * fraction of all that respond to second | | .1023 |

**Supplemental Table 4. Cell lines created in this work**

| Identifier | Name |
|---|---|
| NHM200 | RAW 264.7 Tigre pEF1alpha-NLS-iRFP |
| NHM201 | RAW 264.7 Tigre pEF1alpha-NLS-iRFP pIRF1-IRF1-SYFP2 |
| NHM202 | RAW 264.7 Tigre pEF1alpha-NLS-iRFP pIRF1-IRF1-SYFP2 pCXCL10-NLS-mCerulean-T2A-CXCL10 |
| NHM203 | RAW 264.7 Tigre pEF1alpha-NLS-iRFP pIRF1-IRF1-SYFP2 pCXCL10-NLS-mCerulean-T2A-CXCL10 pCXCL9-CXCL9-T2A-NLS-mCherry |

All cell lines are available upon request from the Hao lab, and all tags are endogenous.
